# A naturally DNase-free CRISPR-Cas12c enzyme silences gene expression

**DOI:** 10.1101/2021.12.06.471469

**Authors:** Carolyn J. Huang, Benjamin A. Adler, Jennifer A. Doudna

## Abstract

Used widely for genome editing in human cells, plants and animals, CRISPR-Cas enzymes including Cas9 and Cas12 provide RNA-guided immunity to microbes by targeting foreign DNA sequences for cleavage. We show here that the native activity of CRISPR-Cas12c protects bacteria from phage infection by binding to DNA targets without cleaving them, revealing that antiviral interference can be accomplished without chemical attack on the invader or general metabolic disruption in the host. Biochemical experiments demonstrate that Cas12c is a site-specific ribonuclease capable of generating mature CRISPR RNAs (crRNAs) from precursor transcripts. Furthermore, we find that crRNA maturation is essential for Cas12c-mediated DNA targeting. Surprisingly, however, these crRNAs direct double-stranded DNA binding by Cas12c using a mechanism that precludes DNA cutting. Cas12c’s RNA-guided DNA binding activity enables robust transcriptional repression of fluorescent reporter proteins in cells. Furthermore, this naturally DNase-free Cas12c enzyme can protect bacteria from lytic bacteriophage infection when targeting an essential phage gene. Together these results show that Cas12c employs targeted DNA binding to provide anti-viral immunity in bacteria, providing a native DNase-free pathway for transient antiviral immunity.

## INTRODUCTION

CRISPR-Cas (<u>c</u>lustered regularly interspaced <u>s</u>hort <u>p</u>alindromic repeats, <u>C</u>RISPR <u>as</u>sociated) systems provide adaptive immunity in bacteria (Barrangou et al., 2007), and also function as powerful tools for genome editing in plants, animals and microbes (Knott and Doudna, 2018). Fundamental to these systems are RNA-guided nucleases, including the Cas9 (type II) and Cas12 (type V) enzyme families that use CRISPR RNA (crRNA) to recognize foreign double-stranded DNA (dsDNA) by forming an R-loop structure in which ∼20 nucleotides of the crRNA (the crRNA “spacer”) base pair with the target strand (TS) of the target DNA, displacing the non-target strand (NTS) (Makarova et al., 2020; Swarts and Jinek, 2018). These Cas proteins also recognize a protospacer-adjacent motif (PAM), a short DNA sequence next to the crRNA-complementary sequence, to prevent autoimmunity and initiate R-loop formation for DNA interference (Gleditzsch et al., 2019).

Multiple studies identified a family of RuvC-containing CRISPR-associated proteins, termed Cas12, whose variants exhibit diverse biochemical and cellular activities (Burstein et al., 2017; Shmakov et al., 2015; Yan et al., 2019). Among them, Cas12c (also known as C2c3), originally found in small DNA fragments from marine and gut metagenomes (Shmakov et al., 2015), shares certain features with other DNA-cutting Cas12 enzymes, but several aspects of this protein are enigmatic. First, although mature guide RNA production is known to be catalyzed by Cas12c with the requirement of a trans-activating crRNA (tracrRNA), the mechanism of RNA recognition and processing is not known (Harrington et al., 2020). Second, despite the demonstration of its gene targeting activity in *E. coli* and the presence of a RuvC catalytic domain with conserved active site residues (Yan et al., 2019), no enzymatic DNA cleavage activity was observed *in vitro* (Harrington et al., 2020). In addition, the lack of detectable DNase activity raised the question of whether or how Cas12c provides bacteria with antiviral protection.

We hypothesized that Cas12c might operate in bacteria using a DNase-free mechanism that is distinct from other Cas12 family members. To test this idea, we first confirmed the inability of the RuvC active site to catalyze DNA cleavage and then tested its activity as a ribonuclease, revealing its function in site-specific precursor crRNA (pre-crRNA) processing. *In vitro* binding assays indicated that despite the lack of target DNA cleavage, Cas12c binds to target DNA through a canonical crRNA-mediated interaction. Cell-based assays in bacteria demonstrated robust repression of transcription by Cas12c without target-activated cleavage. In addition, Cas12c and crRNAs that recognize an essential phage gene protected bacteria from lytic bacteriophage infection. These findings show that a DNA binding-only mechanism is capable of conferring immunity during viral attack. We conclude that Cas12c is a naturally occurring example of an RNA-guided DNA binding enzyme that protects bacteria from infection by target binding rather than target degradation. Furthermore, its immune function is likely to be dependent on RuvC-mediated pre-crRNA processing in its natural host. These results suggest that other functional DNA-targeting CRISPR immune systems may lack enzymatic machinery, and that transcriptional repression may play a larger role in adaptive immunity than previously thought.

## RESULTS

### Cas12c processes its pre-crRNA but cannot cleave DNA *in vitro*

To confirm the repertoire of enzymatic activities that Cas12c has at its disposal for immune function, we began by reconstituting the previously reported pre-crRNA processing activity of Cas12c and probed it in greater detail. Unlike all other auto-processing Cas enzymes (Behler and Hess, 2020; East-Seletsky et al., 2016; Fonfara et al., 2016; Pausch et al., 2020; Yan et al., 2019; Zhang et al., 2018), which cleave the pre-crRNA proximal to or within the repeat sequence that is directly bound by the protein, Cas12c cleaves its pre-crRNA 17 nucleotides (nt) downstream (3’) of the protein-bound repeat (Harrington et al., 2020) (Fig. 1A). We confirmed the previously reported cleavage site, defined by a 17-nt distance downstream of the protein-bound repeat, and showed that it is precise and robust to adjustments in sequence or length on either the 5’ or 3’ end of the pre-crRNA substrate (Fig. S1).

**Figure 1.**
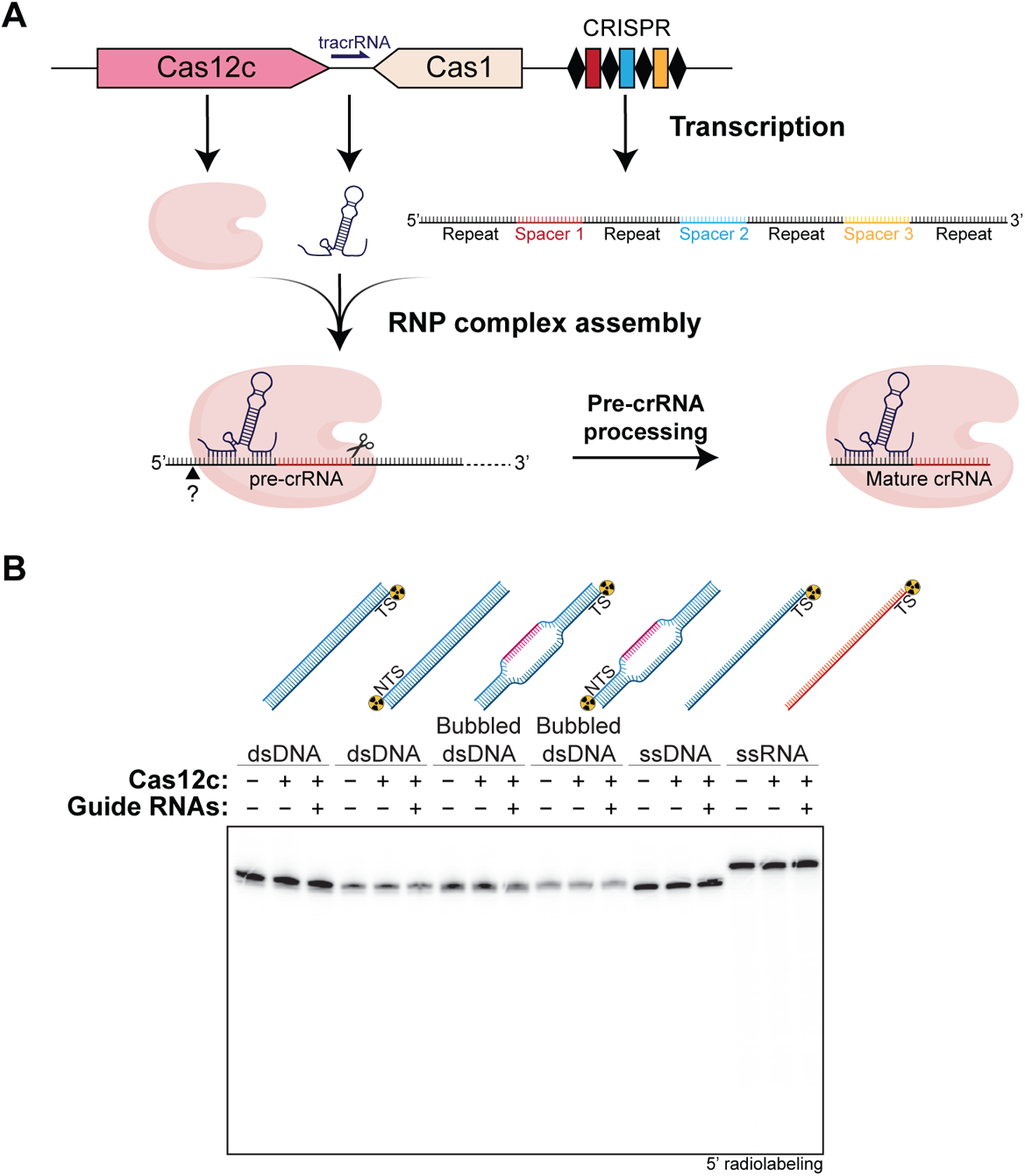
Cas12c processes its pre-crRNA but cannot cleave DNA *in vitro*. (A) Diagram of type V-C CRISPR-Cas12c genomic locus and 3’ pre-crRNA processing by Cas12c. The upstream repeat is processed at the 5’ end by unknown enzymes, as indicated by the question mark. (B) DNA cleavage assay targeting dsDNA, partially base-paired (“bubbled”) dsDNA (dsDNA containing 17 DNA-DNA base pair mismatches throughout the putative region of RNA complementarity), ssDNA, and ssRNA. TS = target strand. NTS = non-target strand.

Next, we tested the ability of Cas12c to perform target-activated DNA cleavage. We tested a variety of different target substrates, including double-stranded DNA, single-stranded DNA, partially base-paired (“bubbled”) DNA and single-stranded RNA. Cas12c failed to cleave these guide-RNA-complementary substrates under all conditions tested (Fig. 1B). Additionally, although non-specific DNA or RNA *trans*-cleavage activity is a property of other Cas12-family enzymes (Chen et al., 2018; Harrington et al., 2018, 2020; Pausch et al., 2020; Wang and Zhong, 2021; Yan et al., 2019), we did not detect such target-activated *trans* cleavage under the conditions tested. These results led us to conclude that Cas12c is a ribonuclease but not a DNase.

### The Cas12c RuvC domain is responsible for pre-crRNA processing

We hypothesized that the RuvC nuclease domain of Cas12c, instead of cleaving DNA, catalyzes pre-crRNA processing. Consistent with this hypothesis, mutation of a RuvC metal-coordinating carboxylate (D928A; described in Yan et al., 2019) abolished Cas12c’s pre-crRNA-processing activity (Fig. 2A). To further test the role of the RuvC domain in pre-crRNA processing, we asked whether the chemical requirements and products of Cas12c’s cleavage activity are consistent with known enzymatic features of RuvC function.

**Figure 2.**
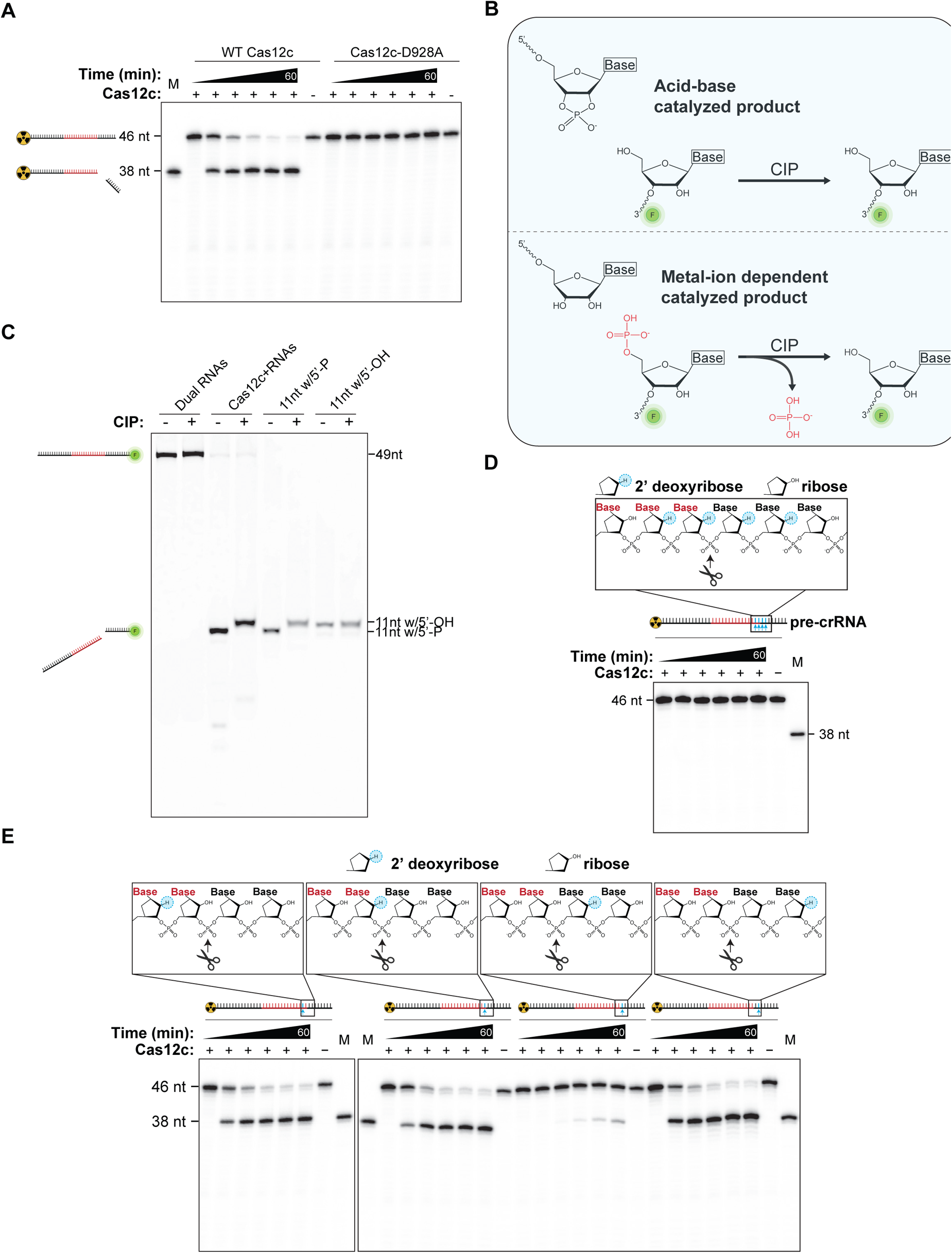
The Cas12c RuvC domain processes pre-crRNA. (A) Pre-crRNA processing assay for Cas12c in dependence of RuvC active site residue variation (D928A). (B) Schematics of the termini chemistry after cleavage occurs by a metal-dependent mechanism and by acid-base catalysis, and respective outcomes of CIP-phosphatase treatment on the 3’ cleavage product. (C) CIP phosphatase treatment identifies the 3’ crRNA cleavage product to possess a 5’ phosphate terminus consistent with metal ion-dependent cleavage. Customized oligonucleotides of the same 11-nt sequence with a 5’-phosphate or a 5’-hydroxyl terminus served as positive and negative controls, respectively. (D) Cas12c was not able to process a DNA/RNA hybrid pre-crRNA substrate containing four 2’-deoxynucleotides spanning the pre-crRNA processing site. The position corresponding to the scissile phosphate is indicated with a scissor icon. (E) Pre-crRNA processing assays with modified pre-crRNAs containing a 2’-deoxynucleotide at indicated positions revealed a requirement of a 2’-hydroxyl group directly downstream of the scissile phosphate for pre-crRNA cleavage.

Because RuvC belongs to a group of enzymes that require divalent cations for activity (Yang and Steitz, 1995), we first investigated the metal-ion dependency of the Cas12c-catalyzed pre-crRNA processing reaction by performing the pre-crRNA processing reaction in the presence of the divalent metal ion chelating reagent EDTA. Results showed that Cas12c cannot process pre-crRNA without access to divalent cations (Fig. S2A). However, further experimentation revealed that Cas12c also cannot bind to pre-crRNA in the presence of EDTA (Fig. S2B), suggesting that Mg^2+^ is likely required for ribonucleoprotein (RNP) complex formation. Therefore, metal ion dependency is not a reliable indicator of the mechanism of catalysis in this case.

We next evaluated the nature of the RNA products generated by Cas12c-catalyzed pre-crRNA cleavage. General acid-base-catalyzed RNA cleavage employs the 2’-hydroxyl group upstream of the scissile phosphate as the attacking nucleophile, producing cleavage products with 2’,3’-cyclic phosphate and 5’-OH termini, respectively (Knott et al., 2017; Swarts et al., 2017) (Fig. 2B). In metal-ion dependent RNA cleavage reactions, water serves as the attacking nucleophile and the 3’ cleavage product has a 5’ phosphate terminus (Pausch et al., 2020) (Fig. 2B). To identify the 5’-terminal chemistry of the Cas12c-generated 3’ cleavage product, we treated the mature crRNA with phosphatase and resolved the product by denaturing polyacrylamide gel electrophoresis (PAGE). If a 5’-phosphate is present in the Cas12c-generated 3’ cleavage product, phosphatase-catalyzed phosphate removal would be expected to reduce the electrophoretic mobility of the treated RNA. Conversely, the lack of a phosphatase-dependent mobility shift would indicate that no removable phosphate was present on the Cas12c-generated 3’ cleavage product. We observed reduced electrophoretic mobility of the 3’ RNA cleavage fragment after CIP treatment (Fig. 2C), consistent with a metal-ion-dependent catalytic mechanism and suggesting that the metal-ion-dependent RuvC domain of Cas12c is responsible for pre-crRNA processing.

Given the lack of detectable *cis* or *trans* DNA cleavage under tested conditions, we further investigated the ability of the RuvC domain to cleave DNA by using a pre-crRNA substrate containing four 2’-deoxynucleotides spanning the pre-crRNA processing site (Fig. 2D). This substrate was not cleavable by Cas12c, indicating that despite its ribonucleolytic activity, Cas12c’s RuvC domain cannot cleave DNA even when presented in the context of the established pre-crRNA substrate. We further demonstrated that only the 2’-hydroxyl group downstream of the scissile phosphate is required for pre-crRNA cleavage (Fig. 2E). Notably, the upstream 2’-hydroxyl group, which is necessary for acid-base catalysis, is not required for pre-crRNA processing by Cas12c (Fig. 2E), consistent with a metal-ion dependent mechanism. Together, our data suggest that the role of the RuvC domain in Cas12c is only for pre-crRNA processing.

### Cas12c catalyzes maturation of an engineered single-guide RNA

Since Cas12c is the first tracrRNA-requiring Cas protein found to catalyze autonomous pre-crRNA processing (Chylinski et al., 2013; Deltcheva et al., 2011; Fonfara et al., 2016; Pausch et al., 2020; Yan et al., 2019), we wondered whether Cas12c’s dual guide RNAs (crRNA and tracrRNA) can be engineered as a single guide RNA (sgRNA) without disrupting Cas12c-catalyzed pre-crRNA maturation. Based on the predicted secondary structure of the mature dual RNAs (Yan et al., 2019) (Fig. 3A), we tested two different pre-sgRNA designs. In the first design, the 3’ end of the tracrRNA was connected by a 5’-GAAA-3’ linker sequence to the unprocessed 5’ end of the crRNA (pre-sgRNA 1) (Fig. 3A). In the second design, the 3’ end of the tracrRNA was instead connected by the same linker to the processed 5’ end of the crRNA (pre-sgRNA 2). We found that Cas12c processes pre-sgRNA 1 as efficiently as when the dual guide RNAs are not fused together (Fig. 3B, C), while pre-sgRNA 2 is processed less efficiently (Fig. 3B, C). The pre-sgRNA design 1 may be useful for future applications of Cas12c.

**Figure 3.**
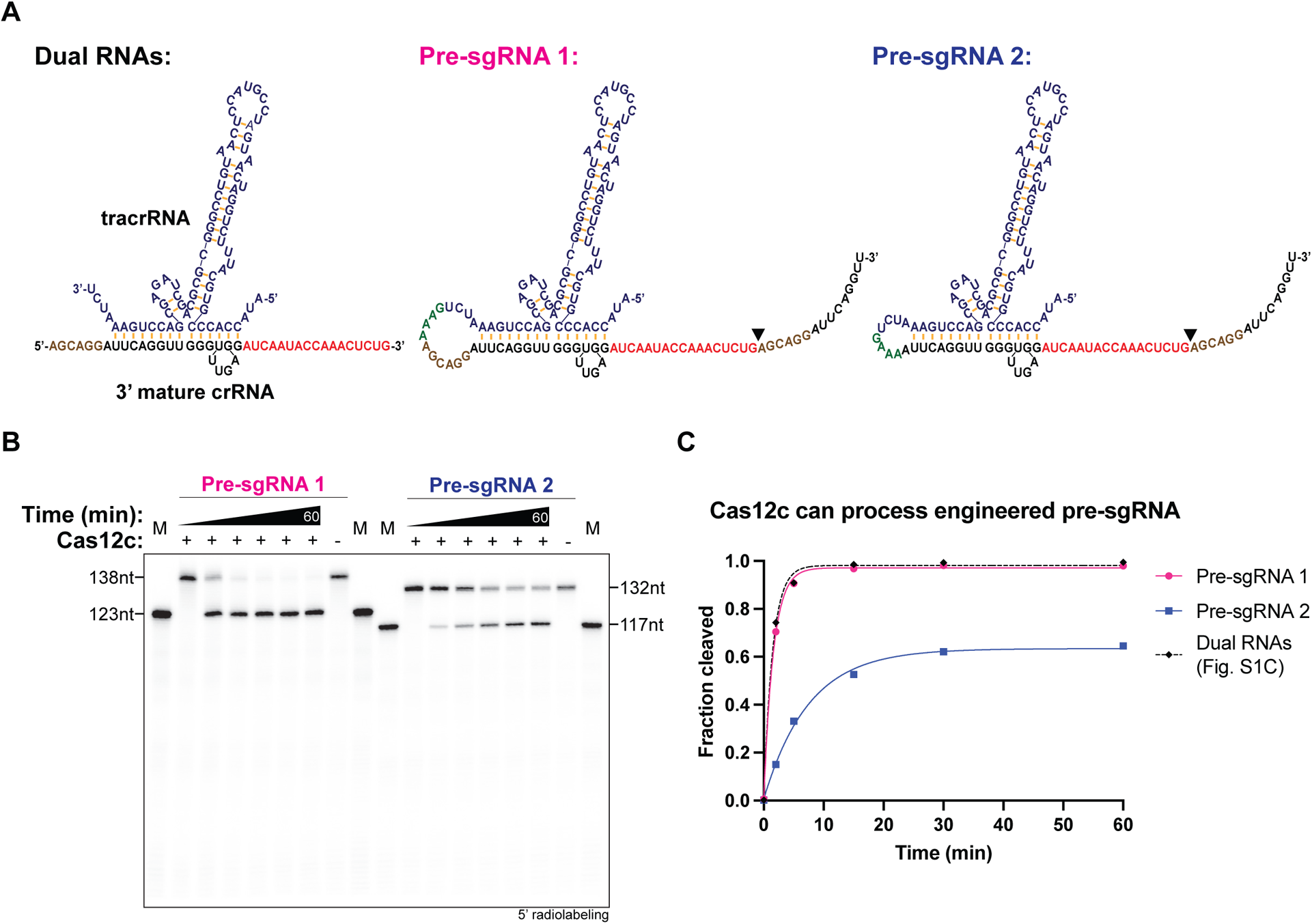
Design and function of pre-sgRNAs for Cas12c. (A) Schematic representation of dual guide RNA (adapted from Yan et al., 2019), pre-sgRNA 1, and pre-sgRNA 2. Spacer sequences are shown in red. The two pre-sgRNA designs differ by the inclusion or exclusion of the 6 nucleotides (shown in brown) present at the 5’ end of the upstream repeat sequence (shown in black). These 6 nucleotides are shown to be processed *in vivo* by RNAseq data (Yan et al., 2019). The cleavage sites by Cas12c in pre-sgRNAs are represented by black triangles. (B) RNA processing assay testing whether Cas12c can process the 2 different pre-sgRNAs. (C) Pre-sgRNA cleavage efficiency compared with pre-crRNA cleavage from Fig. S1C.

### RNA-guided DNA binding by Cas12c depends on pre-crRNA processing

Having demonstrated that Cas12c does not perform target DNA cleavage (Fig. 1B), we next asked whether Cas12c binds guide-RNA-complementary substrates. Filter binding assays indicated that Cas12c binds target dsDNA with a K_D_ in the low nanomolar range and has even stronger affinity for partially base-paired (“bubbled”) DNA (Fig. 4A). These data suggest that like other Cas12 enzymes and Cas9, Cas12c forms an R-loop with complementary dsDNA (Cofsky et al., 2020; Jiang et al., 2016; Lim et al., 2016; Liu et al., 2019; Pausch et al., 2021; Stella et al., 2017; Swarts and Jinek, 2019; Takeda et al., 2021; Yang et al., 2016; Zhang et al., 2021). Cas12c also has moderate binding affinity for target ssDNA (∼26 nM) but does not bind detectably to ssRNA (Fig. 4A). Filter binding assays additionally demonstrated that target DNA binding is sequence-specific and PAM-dependent (Fig. 4A).

**Figure 4.**
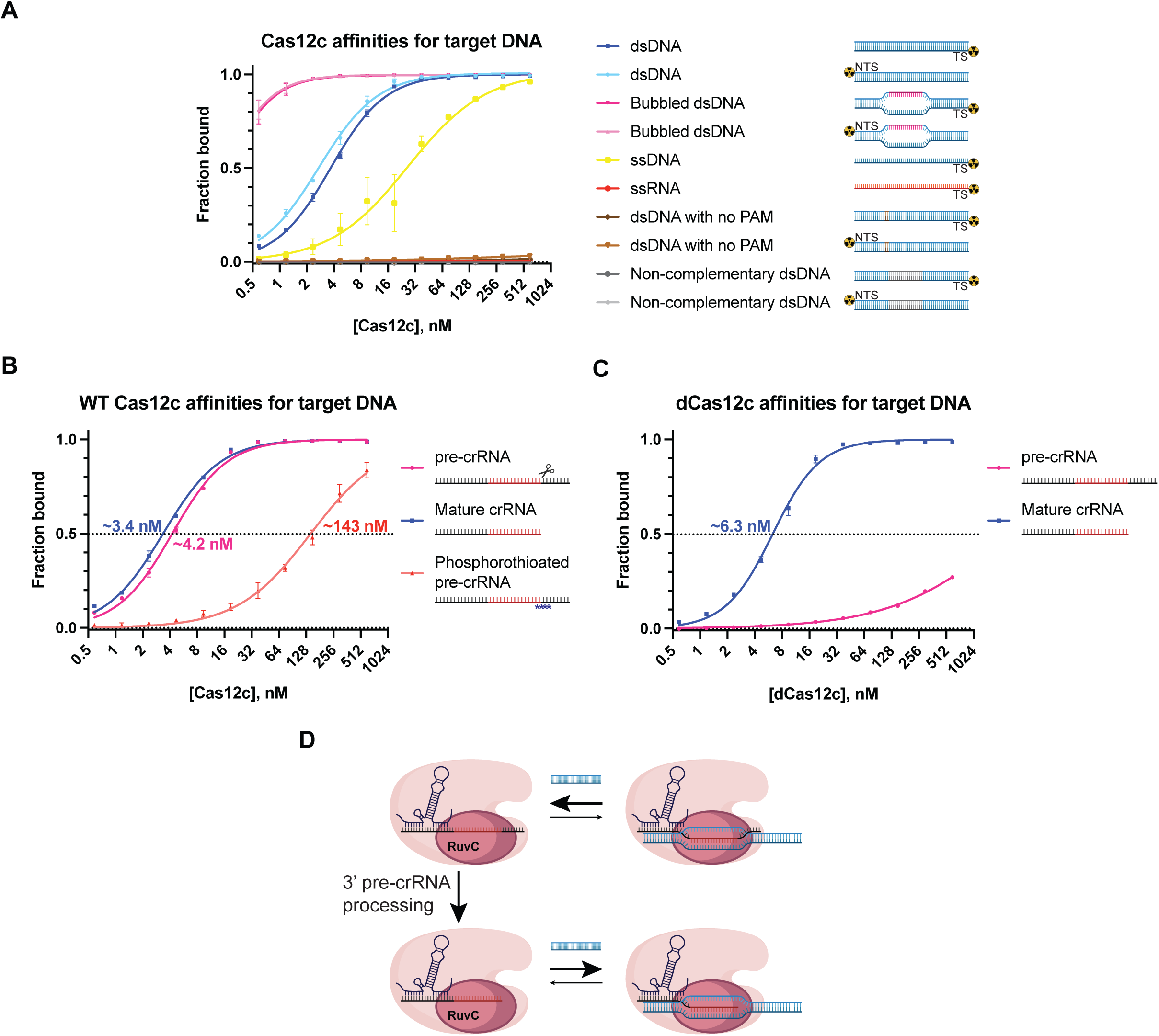
Tight RNA-guided DNA binding by Cas12c depends on pre-crRNA processing. (A) Target DNA binding is sequence-specific and PAM-dependent. Data are from nitrocellulose filter binding assays with radiolabeled dsDNA, partially base-paired (“bubbled”) dsDNA, ssDNA, and ssRNA as a function of Cas12c protein concentration when supplied with a fixed concentration of dual RNAs. K_D_ for fully duplexed dsDNA was 2.7 nM (n=2, 95% CI: 2.6 to 2.9) when NTS was radiolabeled and 3.7 nM (n=2, 95% CI: 3.5 to 3.8) when TS was radiolabeled. K_D_ for bubbled dsDNA was 0.24 nM (n=2, 95% CI: 0.18 to 0.30) when NTS was radiolabeled and 0.25 nM (n=2, 95% CI: 0.19 to 0.32) when TS was radiolabeled. K_D_ for ssDNA was 26.5 nM (n=2, 95% CI: 19.4 to 39.8). (B) The dependence of dsDNA binding by Cas12c on pre-crRNA processing. Data are from filter binding assays with radiolabeled dsDNA as a function of wild-type Cas12c protein concentration in the presence of tracrRNA and pre-crRNA, mature crRNA, or phosphorothioated pre-crRNA. Estimated K_D_ values are reported as shown (n=2). (C) Mature crRNA restores the ability of dead Cas12c (D928A) to bind dsDNA. Data are from filter binding assays with radiolabeled dsDNA as a function of dCas12c protein concentration in the presence of tracrRNA and either pre-crRNA or mature crRNA. Estimated K_D_ values are reported as shown (n=3). (D) Schematic of the proposed model for efficient target dsDNA binding by Cas12c in dependence of pre-crRNA processing.

The pre-crRNA processing activity of Cas12c suggested that crRNA maturation may be essential for target dsDNA binding. To explore this idea, we tested Cas12c’s DNA binding affinity when guided by either pre-crRNA or mature crRNA. Wild-type Cas12c was able to bind to target dsDNA with comparable affinities in each case (Fig. 4B). However, when the pre-crRNA contained phosphorothioates that prevent Cas12c-catalyzed processing (Fig. S1C), Cas12c bound target dsDNA much more weakly (Fig. 4B). Consistent with this observation, we found that Cas12c containing a catalytically deactivating mutation (dCas12c-D928A) was impaired for RNA-guided dsDNA binding when a pre-crRNA was used and behaved similarly to the wild-type Cas12c when a mature crRNA was used (Fig. 4C). These results suggest that pre-crRNA processing is required for the function of the CRISPR-Cas12c system and that without pre-crRNA processing, Cas12c cannot bind target dsDNA efficiently (Fig. 4D). Together these findings imply that the functional role of the RuvC domain in this CRISPR system is distinct from its role in most other systems.

### Cas12c blocks gene expression in cells without invoking target-activated DNase activity

Cas12c can target genes essential for survival in bacteria as detected by depletion of gene-targeting members of a crRNA library transformed into Cas12c-expressing *E. coli* (Yan et al., 2019). This observation could result from Cas12c-mediated cleavage of genomic targets *in vivo*, potentially with the help of *E. coli* host factors, or it could reflect Cas12c-mediated transcriptional silencing through targeted polymerase blockage. To distinguish between these possibilities, we developed a dual-color fluorescence interference assay using a strain of *E. coli* (Qi et al., 2013) in which reporter genes encoding red fluorescent protein (RFP) or green fluorescent protein (GFP) are integrated into the *E. coli* genome. Cas12c-directed cleavage of the chromosomally integrated RFP or GFP would be expected to be toxic to *E. coli* (Fig. 5A) due to chromosome breaks (Cui and Bikard, 2016). In contrast, if Cas12c binds but does not cut target DNA, we expect to observe a loss of the genetically targeted fluorophore without cell death (Fig. 5A), similar to the effect of CRISPRi with a catalytically inactivated Cas9 or Cas12a (Kim et al., 2017; Qi et al., 2013; Zhang et al., 2017).

**Figure 5.**
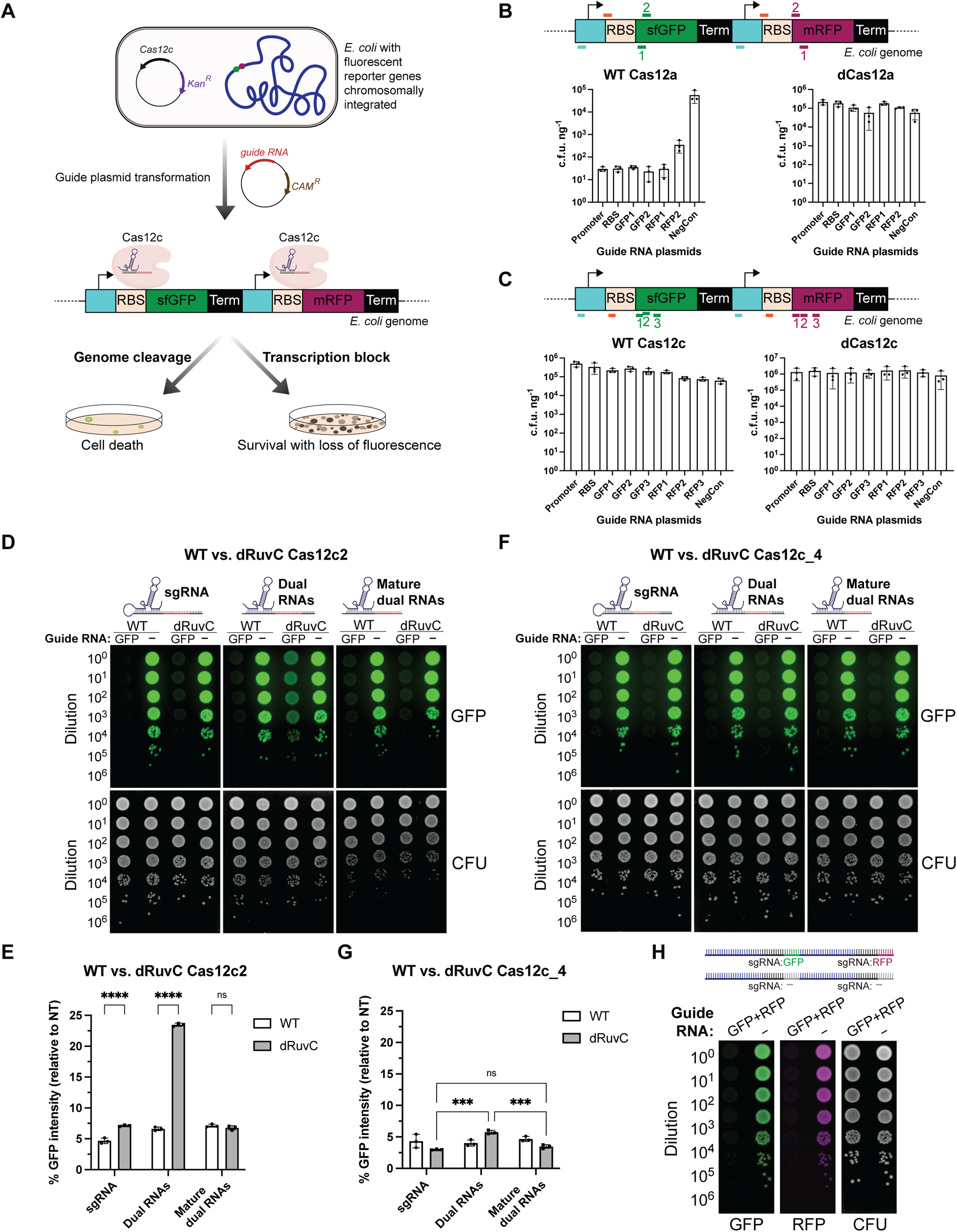
Cas12c blocks gene expression *in vivo* without target-activated DNase activity. (A) Graphical representation of the dual-color fluorescence interference assay used to distinguish target DNA binding from target DNA cleavage *in vivo*. (B) Control fluorescence interference experiments with Cas12a. Wild-type Cas12a with targeting crRNA killed cells, while dead Cas12a with targeting crRNA had no effect in survival but silenced gene expression (see Fig. S3A), as compared to non-targeting negative controls (NegCon). The DNA regions targeted by different spacers (colored lines) used to repress the fluorescent reporters are indicated. c.f.u. ng^-1^ = colony forming unit per nanogram of guide plasmids used in transformation. Data are represented as mean ± SD (n=3). (C) Fluorescence interference experiment with Cas12c. Both wild-type Cas12c and dead Cas12c with targeting guides silenced GFP or RFP expression (Fig. S3B) without killing the cells, as compared to non-targeting negative controls (NegCon). Data are represented as mean ± SD (n=3). (D) Images of the fluorescence interference experiment with the wild-type and RuvC-dead (dRuvC) Cas12c2 (a variant from Yan et al., 2019) when GFP-targeting or non-targeting (–) sgRNA, dual RNAs, or mature dual RNAs were used. (E) Quantification of GFP intensities in targeting guide expressing cells relative to non-targeting guide expressing cells (from Fig. 5D). Data are represented as mean ± SD (n=3). (F) Images of the fluorescence interference experiment with the wild-type and RuvC-dead Cas12c_4 (the focus of this study) when GFP-targeting or non-targeting (–) sgRNA, dual RNAs, or mature dual RNAs were used. (G) Quantification of GFP intensities in targeting guide expressing cells relative to non-targeting guide expressing cells (from Fig. 5F). Data are represented as mean ± SD (n=3). (H) Cas12c can use sgRNAs targeting GFP and RFP coexpressed in a single transcript for multiplexed targeting. Two non-targeting (–) spacers were in place of the targeting spacers as a negative control. Images shown are representative of the effect seen in replicates (n = 3).

We used wild-type LbCas12a (WT Cas12a) and catalytically deactivated LbCas12a (dCas12a) as controls, which exhibited the expected behavior: expression of WT Cas12a killed the bacteria, while dCas12a silenced gene expression without cell death (Fig. 5B, S3A). In contrast, both WT and dCas12c, when guided by a GFP- or RFP-targeting sgRNA, silenced gene expression without killing cells (Fig. 5C, S3B). These results are consistent with biochemical data showing that Cas12c binds but does not cleave target DNA (Fig. 4).

We next tested whether Cas12c-induced repression of essential gene expression in *E. coli* causes cell death. In control experiments using either WT or catalytically deactivated Cas12a, we found that crRNAs targeting an essential gene cause cell death or growth defects (smaller colonies), although the magnitude of the effects depended on the crRNA and were enhanced by the target-activated nuclease activity of WT Cas12a (Fig. S3C). WT Cas12c, paired with guide RNAs targeting essential genes, produced effects similar to those observed using dCas12a (Fig. S3D). This finding suggests that DNA binding by Cas12c blocks transcription of essential genes, inducing cell death or reduced cell proliferation.

Previously it was reported with a slightly different Cas12c ortholog that WT Cas12c but not dCas12c prevented growth of *E. coli* expressing genome-targeting guide RNAs, leading to the conclusion that Cas12c functions by a genome-cutting mechanism (Yan et al., 2019). Since our Cas12c construct (database identifier: Cas12c_4) varies from the previously tested Cas12c2 (Yan et al., 2019) by seven amino acids, none of which are predicted RuvC active site residues, we also performed the CRISPR interference assay using the Cas12c2 variant. We found that both WT Cas12c2 and dCas12c2, when expressed in cells containing targeting sgRNAs, repress expression of a chromosomally-integrated GFP gene without killing the bacteria (Fig. 5D). This finding supports our conclusion that both Cas12c_4 (focus of this study) and Cas12c2 (Yan et al., 2019) are RNA-guided DNA binding proteins that do not cut DNA.

Notably, we found that expression of tracrRNA and pre-crRNA under separate promoters (Yan et al., 2019), rather than as a single pre-sgRNA transcript, produced a more apparent difference in GFP repression between the WT Cas12c2 and dCas12c2 (Fig. 5D, E). Since both tracrRNA and pre-crRNA are required for binding and processing by Cas12c (Fig. S3E), these observations imply that guide RNA maturation limits Cas12c function in cells expressing the dual transcripts, perhaps due to differences in the assembly kinetics of a tripartite versus bipartite complex. This difference in repression between WT Cas12c2 and dCas12c2 was eliminated when the crRNA was produced using a self-cleaving HDV ribozyme at the end of the pre-crRNA transcript (Fig. 5D, E), suggesting that the WT Cas12c2/dCas12c2 discrepancy reported previously (Yan et al., 2019) was due to a difference in pre-crRNA processing capability rather than target dsDNA cleavage activity. Consistent with this conclusion, Cas12c-independent dual RNA processing enabled more robust dCas12c_4-mediated gene repression that is comparable to the repression observed in the presence of sgRNA (Fig. 5F and 5G). We noticed that Cas12c_4-mediated GFP repression is less sensitive to the dual vs. sgRNA switch than Cas12c2, which may be explained if the 7-amino-acid difference yields a difference in the baseline pre-crRNA-processing capacity of each variant. Regardless of these differences between Cas12c2 and Cas12c_4, our data, together with those presented by Yan et al. 2019, are best explained by a model in which Cas12c does not cleave DNA. In addition, pre-crRNA processing is a prerequisite for efficient target dsDNA binding (Fig. 4C, D).

Exploiting its autonomous guide RNA maturation activity, we also tested the ability of Cas12c to induce multiplexed CRISPR interference (CRISPRi) by co-expressing in a single transcript two pre-sgRNAs that target GFP and RFP, respectively (Fig. 5H). Cell-based results showed that in this experiment, Cas12c robustly repressed both RFP and GFP expression (Fig. 5H). This multiplexing property can be useful as a biotechnological tool, as demonstrated for other CRISPR-Cas enzymes (McCarty et al., 2020). Together, these *in vivo* data suggest that RNA-guided Cas12c binding is sufficient for robust transcriptional repression of targeted gene expression without target-activated DNase activity and can be used for biotechnological purposes.

### Naturally DNase-free Cas12c protects cells from bacteriophage infection

In nature, CRISPR-Cas systems are thought to protect prokaryotes from viral infections based on RNA-guided cleavage of foreign nucleic acid, and DNase activity is the key enzymatic function of all known DNA-targeting CRISPR systems (Nussenzweig and Marraffini, 2020; Shmakov et al., 2017; Watson et al., 2021). Having no detectable target-activated DNase activity (Fig. 1B and 5), can Cas12c provide antiphage immunity based on targeted DNA binding? We tested this possibility using a phage plaque assay in which ten-fold dilutions of lambda (λ) phage were plated on lawns of *E. coli* expressing Cas12c and guide RNAs that either target the essential-for-virulence *cro* gene in the phage λ genome (Johnson et al., 1978) or contain a non-targeting sequence. Cells that express Cas12c and one out of the two tested λ-phage targeting sgRNAs resulted in a 5000-fold reduction of plaques, as compared to a non-targeting control (Fig. 6A, B). The extent of plaque reduction induced by Cas12c is comparable to that observed with dCas12a but slightly less robust than wild-type Cas12a (Fig. 6A, B). Together, these results establish Cas12c as the first demonstrated example of a natural DNA-targeting CRISPR ribonuclease that provides antiviral immunity without target-activated DNase activity.

**Figure 6.**
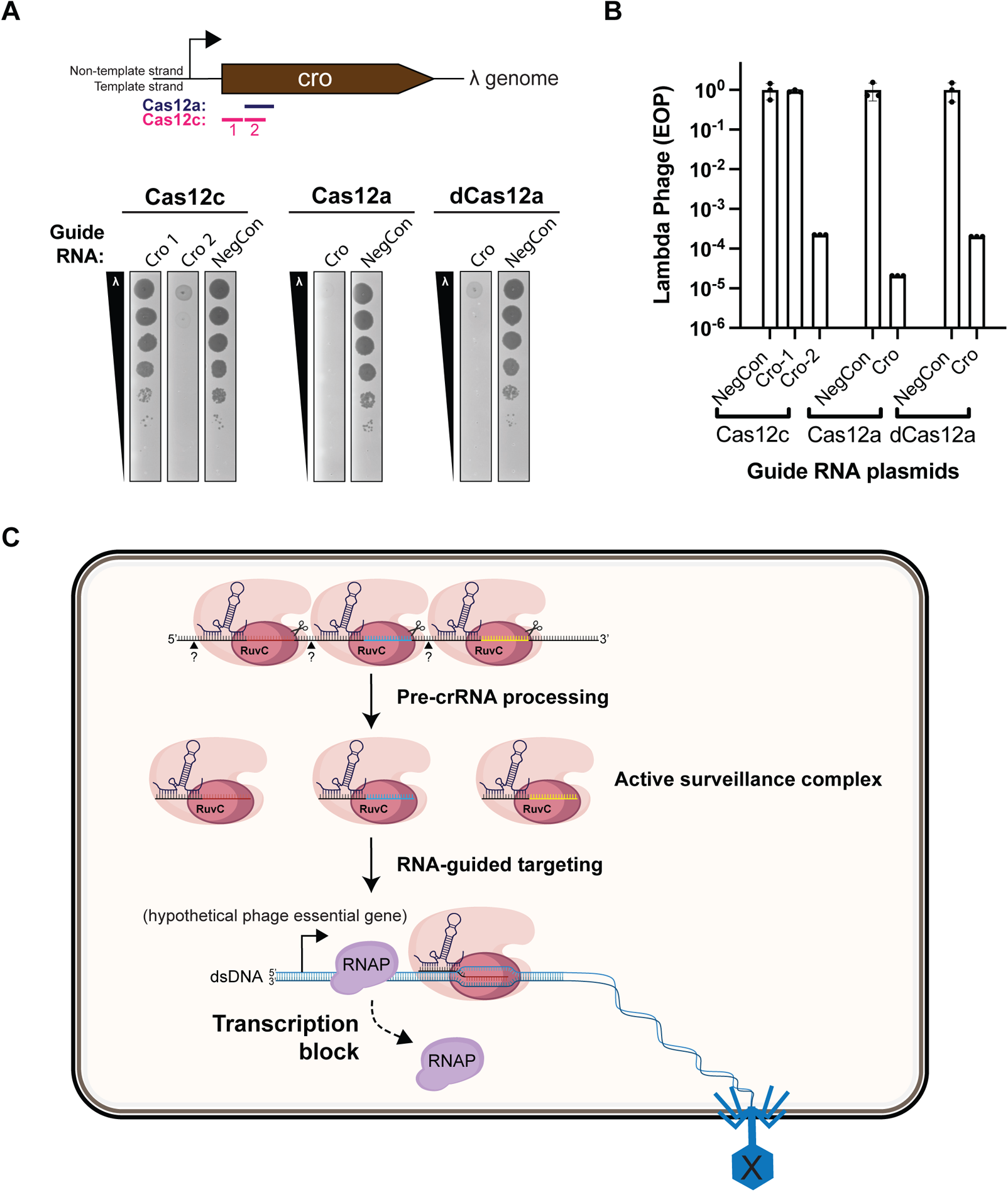
Cas12c protects cells from bacteriophage infection. (A) Phage lambda plaque assay to determine whether Cas12c can confer immunity to phage lambda. Ten-fold serial dilutions of phage lambda were spotted on lawns of *E. coli* strains expressing the specified Cas12 protein and a lambda-targeting guide or a non-targeting guide (as a negative control; NegCon). The DNA regions targeted by different spacers (colored lines) used to repress the *cro* gene in the lambda phage genome are indicated. (B) The efficiency of plaquing (EOP) of lambda phage from Fig. 6A. EOP was calculated as the ratio of plaque forming units (PFUs) on targeting guide expressing cells divided by the number of PFUs on non-targeting guide (NegCon) expressing cells. Data are represented as mean ± SD (n=3). (C) Proposed mechanism of the type V-C CRISPR-Cas12c system. After the CRISPR locus is transcribed into a long precursor transcript, Cas12c, with the help of a tracrRNA, recognizes and binds to pre-crRNA. The long pre-crRNA is cleaved into smaller fragments by Cas12c and unknown enzymes (questionmark), allowing Cas12c to bind target DNA molecules by base-pairing between the crRNA and the DNA target. In this case, Cas12c does not cause double-stranded DNA (dsDNA) breaks and likely only represses gene expression by transcription block. The binding alone is sufficient to confer immunity against certain DNA populations such as essential genes in phages, and this may be Cas12c’s native immune mechanism.

## DISCUSSION

CRISPR-Cas systems have evolved in diverse microbes to provide adaptive immunity against foreign nucleic acids. Until now, immunity provided by the class 2 single-effector type V CRISPR-Cas systems was thought to rely on RNA-guided nuclease activity that targets phage or other foreign molecules, typically at the level of double-stranded DNA cutting. In this study, we show the type V CRISPR effector Cas12c is an RNA-guided DNA-binding enzyme that does not cut target DNA. We found that the RuvC domain of Cas12c, which was previously assumed to cut DNA due to functional comparison to other type V CRISPR enzymes, instead exclusively catalyzes the maturation of pre-crRNA for targeted Cas12c DNA binding. Nonetheless, Cas12c inhibits transcription and can defend bacteria against lytic bacteriophage infection. These results suggest that CRISPR systems can provide anti-phage immunity in the absence of target-directed nuclease activity and, furthermore, that this is Cas12c’s native mechanism (Fig. 6C).

All Cas12-family enzymes contain a RuvC catalytic center, an enzymatic domain resembling Ribonuclease H that catalyzes DNA or RNA phosphodiester cleavage by a “carboxylate-chelated two metal-ion” mechanism (Yang and Steitz, 1995). Enzymes thought to be ancestral to Cas12 including TnpB possess RuvC domain-mediated DNA cutting activity (Altae-Tran et al., 2021; Karvelis et al., 2021), implying that the observed RNA-cutting specificity of Cas12c’s RuvC domain represents a lineage-specific departure from the ancestral state. Notably, the substrate specificities of RuvC domains found within different Cas12 enzymes are divergent. For example, Cas12a’s RuvC domain cleaves DNA only, with a separate active site responsible for pre-crRNA processing (Fonfara et al., 2016). In contrast, the RuvC domain of CasPhi and Cas12g can cleave both DNA and RNA (Pausch et al., 2020; Yan et al., 2019). Based on reported DNA cleavage activity by other Cas12c family members (Wang and Zhong, 2021), the switch to exclusive use of RNA as a substrate by the Cas12c variants tested in this study may have occurred recently.

Repurposing the RuvC active site for exclusive RNA processing suggests that pre-crRNA processing is essential for function. Interestingly, we observed that RuvC-mediated pre-crRNA processing is required for high-affinity DNA binding *in vitro*, but was not as important for transcriptional silencing *in vivo* in a heterologous host. A possible explanation for this observation is that guide RNA expression is under the control of a non-native strong promoter in our assays. If some transcripts undergo abortive transcription or cleavage by host nucleases, the resulting mature guide RNAs may be sufficient to direct unimpeded transcriptional silencing by Cas12c. The preservation of RuvC-catalyzed RNase activity in Cas12c suggests that crRNA maturation is an essential function of Cas12c in its natural host, perhaps due to lower pre-crRNA transcriptional levels or a lack of certain host nucleases.

Prior to this study, single-effector CRISPR-Cas systems that are DNA-targeting yet DNase-free have only been described in the context of transcriptional regulation (Ratner et al., 2019) and transposase association (Strecker et al., 2019) with no indication of immune function. The results presented here suggest that contrary to previous assumptions, immunity against bacteriophage may be achieved solely through RNA-guided DNA binding. Other CRISPR systems for which no immune mechanism has been identified, such as some type IV multi-effector systems (Taylor et al., 2021), may work analogously to Cas12c. The targeting parameters of Cas12c, including mismatch tolerance, seed sequence and strand-dependence, remain to be determined and could be distinct from those of DNA-cleaving Cas12 enzymes. In addition, the structural basis for the substrate specificity swap in Cas12c’s RuvC active site will be fascinating to uncover. Nonetheless, the specific properties of this CRISPR-Cas system, including its minimal PAM requirement (TN), capacity for multiplexed targeting and uniquely precise pre-crRNA processing activity, offer new tool development potential for transcriptional regulation and engineered base editing.

## ACKNOWLEDGMENTS

We thank members of the Doudna lab and the Innovative Genomics Institute for helpful discussions. We would also like to acknowledge Josh Cofsky, Dr. Patrick Pausch, and Dr. Brady F. Cress for advice and guidance. The modified *E. coli* strain for interference assays was kindly provided by Basem Al-Shayeb. This project was funded by grants from the National Science Foundation and by support from the Howard Hughes Medical Institute. CJH acknowledges the support of the T32 NIH Genetics Training Grant (T32GM132022-02). Phage experiments and analyses by BAA were supported by m-CAFEs Microbial Community Analysis & Functional Evaluation in Soils, a project led by Lawrence Berkeley National Laboratory based upon work supported by the U.S. Department of Energy, Office of Science, Office of Biological & Environmental Research. JAD is an Investigator of the Howard Hughes Medical Institute. Plasmids are available from Addgene (addgene.org) or by request.

## AUTHOR CONTRIBUTIONS

Conceptualization: CJH, JAD Investigation: CJH, BAA Methodology: CJH, BAA, JAD Supervision: JAD Writing: CJH, BAA, JAD

## MATERIALS AND METHODS

### Generation of plasmids

For protein purification, Cas12c bacterial expression plasmid (10xHis-MBP-TEVcs-Cas12c) was described in (Harrington et al., 2020) and modified (R965H) to match with the original protein sequence of Cas12c_4 in the JGI IMG database. For *in vivo* experiments, Cas12a and Cas12c genes were cloned into a low-copy plasmid backbone that contains a SC101 origin of replication and KanR. The backbone of guide RNA plasmids was amplified from pBFC0423 described in (Knott et al., 2019) so that sgRNA or crRNA expression are under control of a strong constitutive promoter (proD). Dual guide RNA plasmids were constructed from Integrated DNA Technologies (IDT) gBlocks® so that the expressions of most of the concatenated non-coding RNA sequence (Yan et al., 2019), which includes tracrRNA, and crRNA are under control of separate proD promoters in opposite directions. A list of plasmids is described in Table S1. Plasmid sequences and maps will be made available on addgene.org. To reprogram the guide RNA plasmids for targeting different loci, guide sequences were exchanged via Gibson Assembly or Golden Gate assembly to encode the guide for the selected target site (guide spacer sequences are listed in Table S1B).

### Nucleic acid preparation

DNA oligonucleotides were synthesized commercially (IDT) and purified by denaturing PAGE, ethanol-precipitated, and resuspended in water before being used in cleavage assays. RNA substrates were either synthesized commercially (IDT, Integrated DNA Technologies) or generated through *in vitro* transcription, which was described previously (Cofsky et al., 2020). For *in vitro* T7 RNA polymerase transcription, double-stranded DNA templates were assembled from several overlapping DNA oligonucleotides (IDT) by PCR or by annealing a short oligonucleotide containing the T7 promoter sequence to a long ssDNA oligonucleotide template (IDT). Synthesized or transcribed RNA fragments were PAGE-purified and resuspended in RNA storage buffer (0.1 mM EDTA, 2 mM sodium citrate, pH 6.4). All oligonucleotide identities and sequences are listed in Table S2.

### Protein expression and purification

Cas12c expression and purification were performed as previously described (Chen et al., 2018) with modifications. Briefly, *E. coli* Rosetta (DE3) (Novagen) containing Cas12c expression plasmids were grown in Terrific Broth supplemented with 100 μg/mL ampicillin and 20 μg/mL chloramphenicol to an OD_600_ of 0.4, cooled down on ice for 15 min before induction with 0.5 mM IPTG and overnight growth at 16 °C. Cells were harvested by centrifugation and re-suspended in lysis buffer (50 mM HEPES-Na, pH 7.5, 500 mM NaCl, 20 mM imidazole, 1 mM TCEP, 5% (v/v) glycerol, 1 tablet of cOmplete Protease Inhibitor Cocktail/50 mL, 0.25 mg/mL lysozyme, and 0.5% TritonX-100) and lysed by sonication. After centrifugation, the soluble fraction of the lysate was loaded onto a Ni-NTA Superflow Cartridge (QIAGEN) pre-equilibrated in wash buffer (50 mM HEPES-Na, pH 7.5, 500 mM NaCl, 20 mM imidazole, 1 mM TCEP, 5% (v/v) glycerol). Bound proteins were washed with wash buffer until UV baseline and eluted in elution buffer (50 mM HEPES-Na, pH 7.5, 500 mM NaCl, 300 mM imidazole, 1 mM TCEP, 5% glycerol). The buffer of the eluate was exchanged to ion exchange buffer A (50 mM HEPES-K, pH 7.5, 200 mM KCl, 1 mM TCEP, 5% (v/v) glycerol) using a HiPrep 26/10 Desalting Column (Cytiva) before the addition of TEV protease. After overnight cleavage at 4°C, proteins were loaded into a MBPTrap HP column (Cytiva) connected to a HiTrap Heparin HP Column (Cytiva) and washed until UV baseline. The TEV-cleaved proteins were eluted from the Heparin column with a KCl gradient and concentrated to 2 mL before injection into a HiLoad 16/600 Superdex 200 pg column (Cytiva). The gel filtration buffer contained 20 mM HEPES-K, pH 7.5, 200 mM KCl, 1 mM TCEP, 5% (v/v) glycerol. Peak fractions were verified by SDS-PAGE, and the concentrations of purified proteins were determined using a NanoDrop 8000 Spectrophotometer (Thermo Scientific).

### Pre-crRNA processing assays

Pre-crRNA processing assays were performed as previously described (Harrington et al., 2020). Briefly, the reactions were carried out in cleavage buffer containing 20 mM Tris-Cl (pH 7.5 at 37°C), 150 mM KCl, 5 mM MgCl_2_, and 5% (v/v) glycerol. For radiolabeling experiments, the pre-crRNA substrates were 5′-radiolabeled with T4 PNK (NEB) in the presence of gamma ^32^P-ATP. In a typical pre-crRNA processing reaction, the concentrations of Cas12c, tracrRNA and ^32^P-labeled pre-crRNA substrates were 100 nM, 125 nM and 3 nM, respectively. Prior to the addition of Cas12c to the reaction, tracrRNA and pre-crRNA were pre-annealed in 1X Cleavage Buffer. Reactions were incubated at 37°C, and an aliquot of each reaction was quenched with 2x Quench Buffer (94% (v/v) formamide, 30 mM EDTA, 400 μg/mL heparin, 0.2% SDS, and 0.025% (w/v) bromophenol blue) at 0, 2, 5, 15, 30, and 60 min. RNA hydrolysis ladders were prepared by incubating RNA probes in 1X RNA Alkaline Hydrolysis Buffer containing 50 mM Sodium Carbonate [Na_2_CO_3_], pH 9.2, and 1 mM EDTA at 95°C for 1, 5, and 15 mins before the addition of 2x Quench Buffer. Quenched reactions were incubated at 95°C for 3 min, and products were then resolved by denaturing PAGE (10% acrylamide:bis-acrylamide 19:1, 7 M urea, 1X TBE). Gels were dried (3 hr, 80 °C) on a Model 583 Gel Dryer (Bio-Rad) and exposed to a phosphor screen. Phosphor screens were imaged on an Amersham Typhoon phosphorimager (GE Healthcare). Phosphorimages were quantified using ImageQuant software (GE Healthcare). In pre-sgRNA processing reactions, the concentrations of Cas12c and ^32^P-labeled pre-sgRNA substrates were 100 nM and 3 nM, respectively.

For experiments with 3’ 6-FAM-labeled pre-crRNA, the concentrations of Cas12c, tracrRNA, and 3’ 6-FAM-labeled pre-crRNA substrates were 2000 nM, 2200 nM and 200 nM, respectively. For termini chemistry analysis, 7 μL of the processing reaction products were treated with 10 units Quick CIP (NEB) in 1X CutSmart buffer (NEB) for 1 hr at 37 °C. The 2X Quenching Solution for fluorophore experiments contained 97% formamide and 30 mM EDTA. Products were resolved by 20% denaturing PAGE gel and visualized with Amersham Typhoon Biomolecular Imager (GE Healthcare). Oligonucleotide identities are shown in Table S3.

### Radiolabeled DNA cleavage assays

Radiolabeled DNA cleavage assays were performed as previously described (Harrington et al., 2020). Briefly, the reactions were carried out in the same cleavage buffer described above for pre-crRNA processing. The tested interference substrates of either target strand (TS) or non-target strand (NTS) were 5′-radiolabeled with T4 PNK (NEB) in the presence of gamma ^32^P-ATP. To form dsDNA substrates, the labeled substrate was annealed with excess cold TS or NTS depending on the labeled strand. The final concentrations of Cas12c, tracrRNA, crRNA, and ^32^P-labeled interference substrates were 100 nM, 125 nM, 125 nM and 3 nM, respectively. Before the addition of the labeled interference substrates at 37°C, Cas12c ribonucleoprotein was pre-complexed by incubating with its pre-annealed dual guide RNAs at 37 °C for 5 min, then 25 °C for 25 min. Reactions were incubated for 60 min and quenched with 2x Quench Buffer (described above for pre-crRNA processing). The quenched reactions were heated to 95°C for 5 min. The reaction products were resolved by 10% denaturing PAGE and phosphorimaging. Oligonucleotide identities are shown in Table S3.

### Filter binding assays

The filter binding assays were performed as previously described (Cofsky et al., 2020). Briefly, complexes were formed in 1X binding buffer (20 mM Tris-Cl, pH 7.9 at 25°C, 150 mM KCl, 5 mM MgCl_2_, 1 mM TCEP, 50 µg/mL heparin, 50 µg/mL bovine serum albumin, 5% glycerol). In a typical binary complex binding assay, Cas protein was first diluted down from 600 nM in series in binding buffer, added to a fixed concentration of tracrRNA (750 nM for all protein dilutions), and was incubated with radiolabeled RNA (100 pM) at 37 °C for 5 min, then 25 °C for at least 1 hr. For ternary complex binding assays, Cas protein was first diluted down from 600 nM in series in binding buffer, added to a fixed concentration of guide RNAs (750 nM), and incubated at 37 °C for 5 min, then 25 °C for 25 min. This complex was then added to the radiolabeled DNA probe (100 pM) and incubated at 37 °C for 5 min, then 25 °C for at least 1 hr. HT Tuffryn (Pall), Amersham Protran, and Amersham Hybond-N+ (GE Healthcare) membranes were equilibrated in 1X membrane wash buffer (20 mM Tris-Cl, pH 7.9 at 25°C, 150 mM KCl, 5 mM MgCl_2_, 1 mM TCEP, 5% glycerol) and assembled on a vacuum dot-blot apparatus. The membranes were washed once with 30 µL 1X wash buffer before radioactive samples were applied to the membranes by low vacuum. Membranes were then washed once with 40 µL 1X wash buffer, air-dried, and analyzed by phosphorimaging. Data were quantified with ImageQuant TL Software (GE Healthcare) and fit to a binding isotherm using Prism (GraphPad Software). “Fraction bound” is defined as (background-subtracted volume of Protran spot)/(total background-subtracted volume of Protran spot + Hybond N+ spot). For DNA binding, data were best fit by a model that included an exponent on the concentration terms. The physical basis for this dependency is unknown. Dissociation constants (K_D_) with 95% confidence intervals (CI) and number of independent replicates (n) are reported in the figure legends, when appropriate. For assays testing complex assembly in EDTA-containing buffer, 25 mM EDTA was substituted for 5 mM MgCl_2_. Oligonucleotide identities are shown in Table S3.

### Dual-color fluorescence interference assay

The strain used for all *in vivo* assays in this study was *E. coli* MG1655 with sfGFP and mRFP chromosomally integrated at the *nsf*A locus, originally described by Qi et al., 2013 and modified with the removal of KanR. Guide RNA plasmids were transformed into electrocompetent cells containing the Cas protein using a MicroPulser Electroporator (Bio-Rad). Cells were recovered for 100 min at 37°C in LB broth, and tenfold serial dilutions were plated on Kan+CAM media in Nunc Rectangular Dishes (Thermo Scientific). Plates were incubated at 37°C for 13-17 hours. GFP images were taken on an Amersham Typhoon Biomolecular Imager (GE Healthcare) with a Cy2 525BP20 filter or on a ChemiDoc MP Imaging System (Bio-Rad) with Blue Epi illumination as excitation source and a 530/28 emission filter. RFP images were taken on the Typhoon Imager with a Cy3 570BP20 filter or on the ChemiDoc imager with Green Epi illumination as excitation source and a 605/50 emission filter. Colonies were visualized using the ChemiDoc imager with White Trans illumination as excitation source and a standard filter, and colony forming units (CFUs) were counted. For experiments comparing WT vs. Dead Cas12c transcriptional repression levels, the GFP intensity of undiluted spots was quantified using Image Lab software (Bio-Rad) and divided by the CFU intensity of the same spots. This ratio is normalized by the ratio of the corresponding non-targeting guide sample so that % GFP intensity is 100% for non-targeting guide samples. All assays were performed in triplicate, and independent transformation was performed for each replicate. Plasmid identities are shown in Table S3.

### Bacteriophage plaque assays

Bacteriophage assays were conducted using a modified double agar overlay protocol using phage λ cI857 (Knott et al., 2019). Briefly, *E.coli* (NEB® 10-beta) containing both a Cas effector and gRNA plasmid (Table S3) were grown overnight 37°C, 200 rpm. To perform plaque assays, 100 µL of saturated overnight culture was mixed with molten LB Lennox top agar supplemented with appropriate antibiotics and decanted onto a corresponding LB Lennox Agar plate (to final overlay concentrations of 1% (w/v) tryptone, 0.5% (w/v) yeast extract, 0.5% (w/v) NaCl, 0.7% (w/v) agar, 50 µg/mL kanamycin sulfate, and 34 µg/mL chloramphenicol hydrochloride). This overlay was left to dry for 15 minutes under microbiological flame. 10X Serial dilutions of λ cI857 were performed in SM buffer (Teknova), and 2 µL of each dilution were spotted onto the top agar and allowed to dry for 10 minutes. Plaque assays were incubated at 37°C for 12-16 hours. After overnight incubation, plaques were scanned using a standard photo scanner and plaque forming units (PFUs) enumerated. In cases where individual PFUs were not enumerable, but clearings were observed at high phage concentrations, the most concentrated dilution at which no plaques/clearings were observed was counted as 1 PFU. Efficiency of plaquing (EOP) calculations for a given condition were performed by normalizing the mean of PFU for a condition to the mean PFU of a non-targeting control: mean(PFU_condition_)/mean(PFU_negativecontrol_). All plaque assays were performed in biological triplicate (three experiments carried out on different days using independent bacterial cultures, independently prepared bottom and top agar, and freshly prepared bacteriophage dilutions).

## SUPPLEMENTAL FIGURE LEGENDS

**Supplemental Figure S1.**
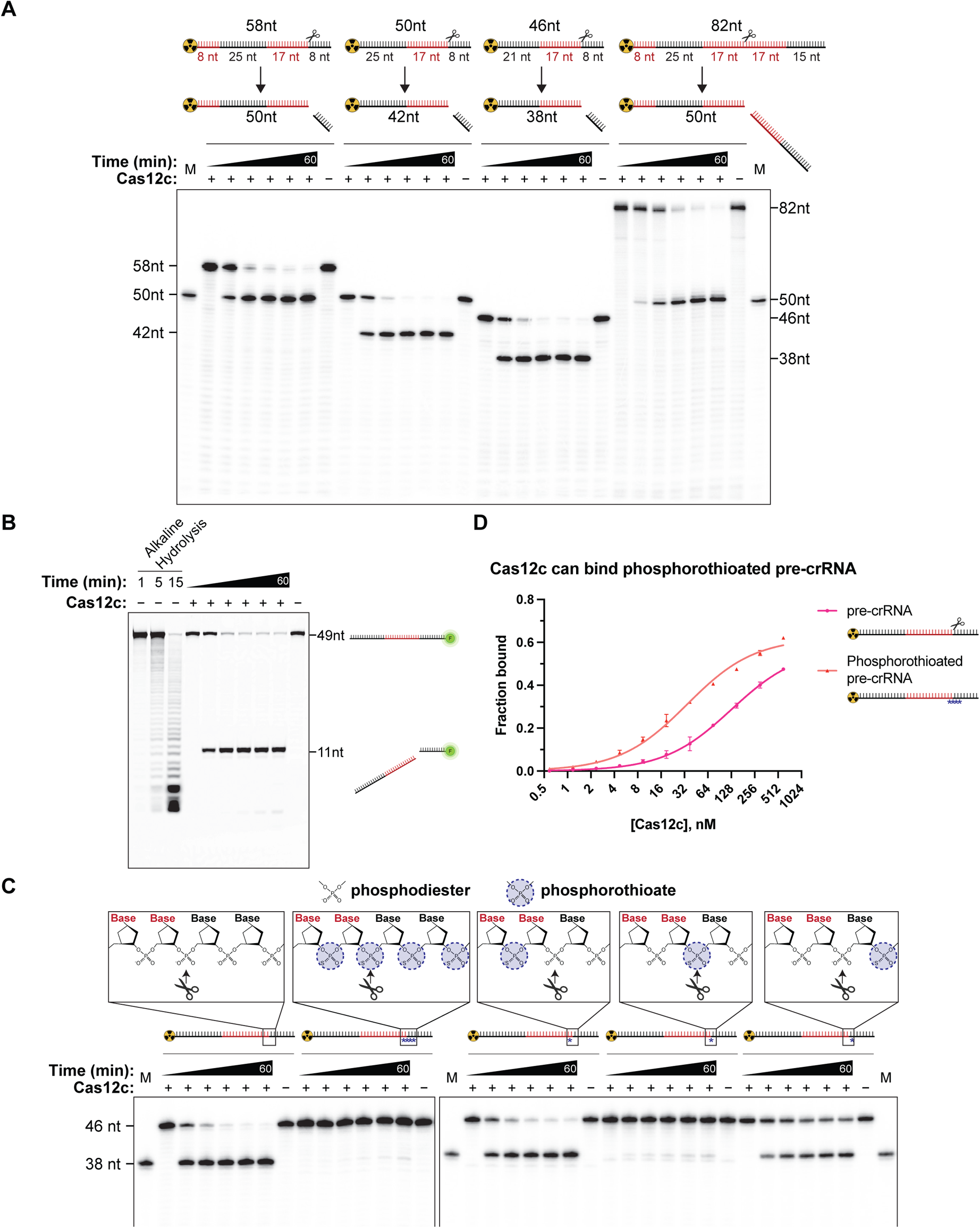
Related to Figure 1; Cas12c pre-crRNA processing is precise and occurs 3’ to the spacer. (A) Time courses of pre-crRNA processing activity of Cas12c when the pre-crRNA has a 5’ extension (this substrate was tested in Harrington et al., 2020), the original length of 5’ (upstream) protein-bound repeat (25 nt), a more processed version of the 5’ repeat sequence, or two spacers in between the upstream and downstream repeat sequences. Repeat sequences are shown in black, and spacer sequences are shown in red. M = Marker. The results confirmed that Cas12c processes pre-crRNA at the expected position, that is 17 nt away from the upstream protein-bound repeat, regardless of the 3’ (downstream) sequence. (B) Time course of RNA processing activity of Cas12c when a pre-crRNA was labeled with 6-FAM fluorophore at the 3’ end to further determine whether Cas12c severs the RNA at one site or more than one site. The result showed that Cas12c-mediated cleavage occurred at the position expected from the 5’-labeled experiments, indicating that Cas12c makes a single precise endonucleolytic cut between the spacer and the repeat. Fig. 2C further confirms that the product here is exactly 11 nt. (C) Phosphorothioate inhibition of RNA processing to test the precision and rigidity of Cas12c-catalyzed RNA cleavage. Different variants of pre-crRNAs are shown, with purple circles indicating phosphorothioate linkages. The position corresponding to the scissile phosphate is indicated with a pair of scissors. M = Marker. A single phosphorothioate modification at the scissile phosphate group almost completely abolished Cas12c processing activity, and the effect was slightly stronger when the flanking linkages were simultaneously phosphorothioated. Note that none of the phosphorothioate modifications caused a shift in the position of cleavage, suggesting that Cas12c-catalyzed pre-crRNA uses a strict and precise ruler mechanism. Modification of the linkage 3’ to the scissile phosphate reduced the extent of processing by ∼40% at 60 mins, suggesting that one of the two non-bridging oxygens may be important for catalysis. (D) Phosphorothioate inhibition of RNA processing was not due to defects in binding. Data are from nitrocellulose filter binding assays with radiolabeled wild-type or phosphorothioated pre-crRNA as a function of Cas12c protein concentration when supplied with a constant concentration of tracrRNA.

**Supplemental Figure S2.**
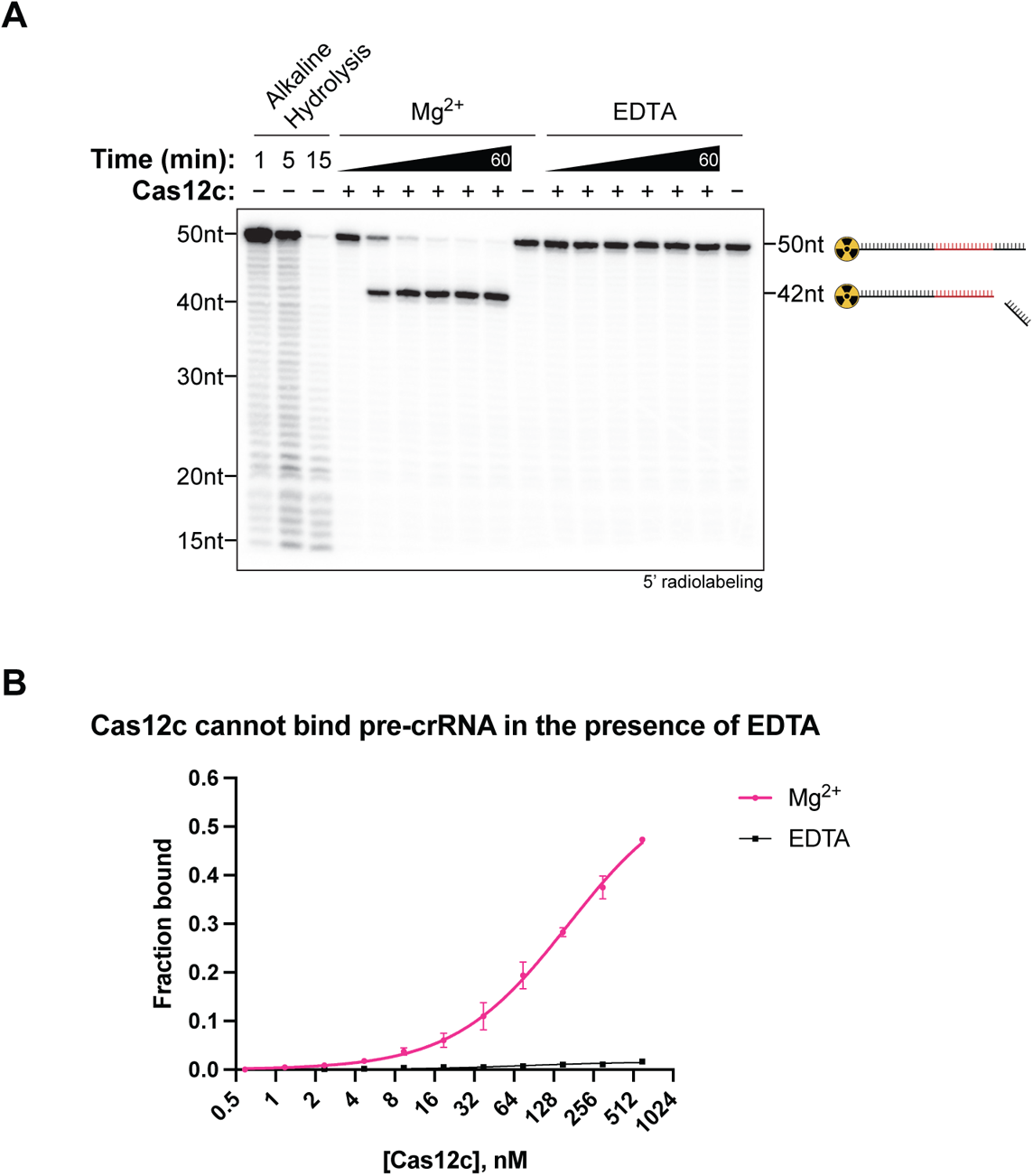
Related to Figure 2; Dependence of pre-crRNA processing and binding on divalent metal ion. (A) EDTA inhibits pre-crRNA processing. Data are from a time course experiment testing the RNA processing activity of Cas12c in the presence of Mg^2+^ or EDTA. (B) EDTA inhibits pre-crRNA binding (binding is metal ion dependent). Data are from nitrocellulose filter binding assays with radiolabeled pre-crRNA as a function of Cas12c protein concentration in the presence of Mg^2+^ or EDTA, when supplied with a constant concentration of tracrRNA.

**Supplemental Figure S3.**
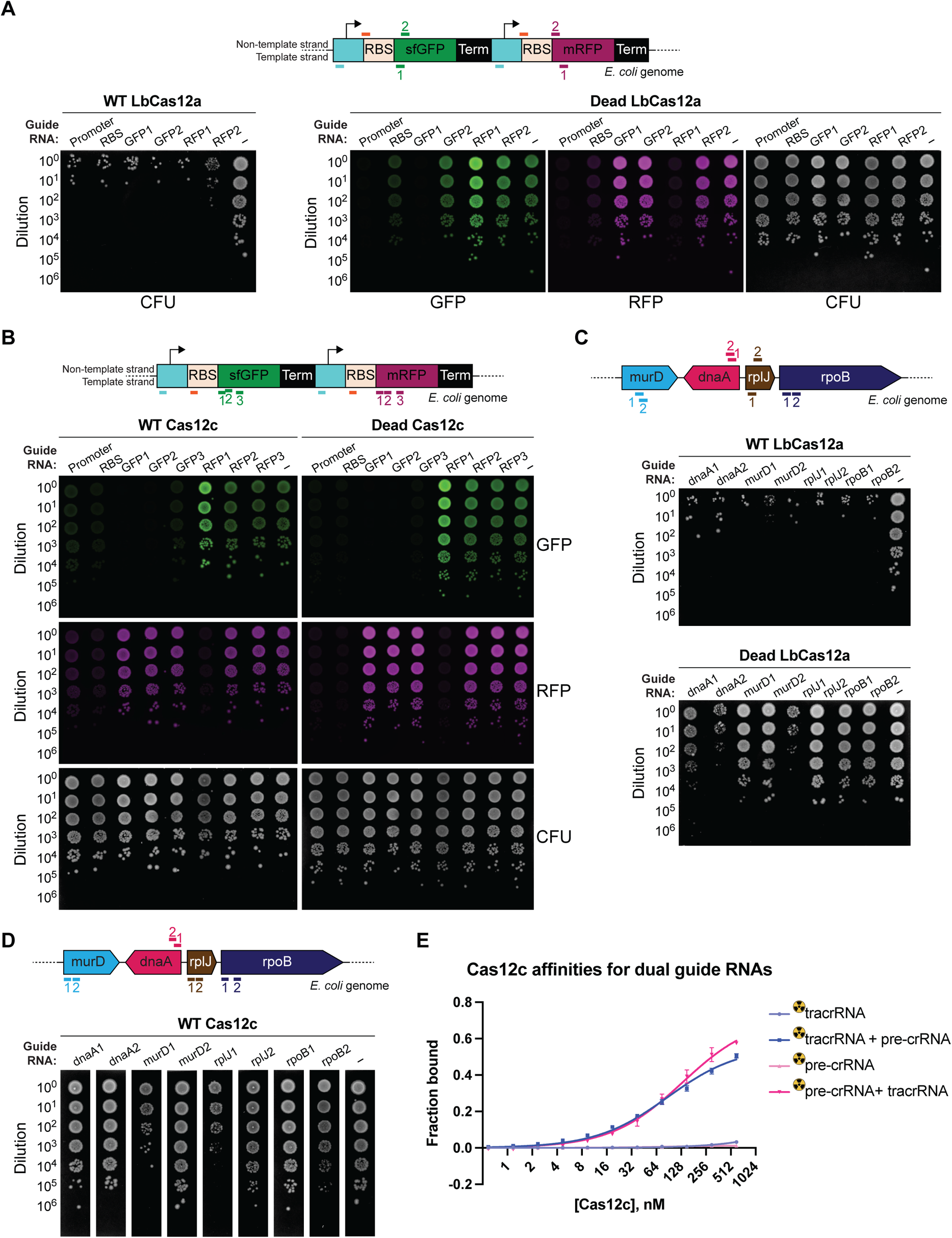
Related to Figure 5; Cas12c blocks gene expression *in vivo* without target-activated DNase activity. (A) Images of fluorescence interference experiments of Cas12a. Wild-type Cas12a with targeting crRNA killed cells, while dead Cas12a with targeting crRNA had no effect in survival but silenced gene expression, as compared to non-targeting (–) controls. It is normal if dead Cas12a with a targeting crRNA produced no effect in gene repression since interference by repression is known to be highly guide-dependent. Images shown are representative of the effect seen in replicates (n = 3). (B) Images of fluorescence interference experiment with Cas12c. Both wild-type Cas12c and dead Cas12c with selected targeting sgRNAs silenced GFP or RFP expression without killing the cells, as compared to non-targeting (–) controls. Images shown are representative of the effect seen in replicates (n = 3). (C) Interference assay to test whether dead Cas12a with crRNAs targeting essential host gene(s) can cause cell death. Data suggest that dead Cas12a with selected crRNAs (targeting dnaA and rplJ_1) could result in depletion of cells containing those guides, as compared to non-targeting (–) controls. WT Cas12a was included for comparison. Images shown are representative of the effect seen in replicates (n = 3). (D) Interference assay to test whether wild-type Cas12c with sgRNA targeting essential host gene(s) can cause cell death. Data showed that wild-type Cas12c with selected crRNAs could cause cell death (murD_1 and rplJ_1) or slower growth (rpoB_2), resulting in depletion of cells containing those guides, as compared to non-targeting (–) controls. Images shown are representative of the effect seen in replicates (n = 3). (E) Cas12c binds to the complex from tracrRNA and crRNA. Data are from filter binding assays with radiolabeled tracrRNA or crRNA as a function of Cas12d protein concentration when they were alone or when two RNAs were present in the binding reaction.

