## Supplemental Tables 1-3 for "A naturally DNase-free CRISPR-Cas12c enzyme silences gene expression"

### Supplementary Table S1: Plasmids

| **Table S1A: Plasmids containing Cas protein** | | | |
|---|---|---|---|
| **Plasmid ID** | **Description** | **Selection marker** | **Addgene number** |
| pCJH002 | Bacterial protein expression plasmid of WT Cas12c. This is the R965H version of the Cas12c in Harrington et al., 2020. | Ampicillin | Yes |
| pCJH003 | Bacterial protein expression plasmid of RuvC-dead mutant (D928A version of pCJH002) | Ampicillin | - |
| pCJH019 | WT Cas12c in a plasmid containing a SC101 origin and *lac* promoter for *in vivo* interference assay | Kanamycin | Yes |
| pCJH020 | RuvC-dead Cas12c (D928A) in a plasmid containing a SC101 origin and *lac* promoter for *in vivo* interference assay | Kanamycin | - |
| pCJH021 | WT Cas12c2 (Yan et al., 2019) in a plasmid containing a SC101 origin and *lac* promoter for *in vivo* interference assay | Kanamycin | Yes |
| pCJH022 | RuvC-dead Cas12c2 (D928A) (Yan et al., 2019) in a plasmid containing a SC101 origin and *lac* promoter for *in vivo* interference assay | Kanamycin | - |
| pCJH027 | WT LbCas12a (Chen et al., 2018) in a plasmid containing a SC101 origin and lac promoter for *in vivo* interference assay | Kanamycin | Yes |
| pCJH028 | Dead LbCas12a (D832A) (Chen et al., 2018) in a plasmid containing a SC101 origin and lac promoter for *in vivo* interference assay | Kanamycin | - |

| **Table S1B: Guide RNA plasmids for *in vivo* assays** | | | | |
|---|---|---|---|---|
| **Plasmid ID** | **Description** | **Selection marker** | **Addgene number** | **Spacer sequence** |
| pCJH007 | Plasmid expressing Cas12c sgRNA targeting the promoter of integrated GFP/RFP | Chloramphenicol | - | ACAGCTAGCTCAGTCCT |
| pCJH008 | Plasmid expressing Cas12c sgRNA targeting the RBS sequence of integrated GFP/RFP | Chloramphenicol | - | AATTCATTAAAGAGGAG |
| pCJH009 | Plasmid expressing Cas12c sgRNA targeting GFP (1) | Chloramphenicol | Yes | TCCCAATTCTTGTTGAA |
| pCJH010 | Plasmid expressing Cas12c sgRNA targeting GFP (2) | Chloramphenicol | - | AATTAGATGGTGATGTT |
| pCJH011 | Plasmid expressing Cas12c sgRNA targeting GFP (3) | Chloramphenicol | - | CACTACTGGAAAACTAC |
| pCJH012 | Plasmid expressing Cas12c sgRNA targeting RFP (1) | Chloramphenicol | - | GCGAGTAGCGAAGACGT |
| pCJH013 | Plasmid expressing Cas12c sgRNA targeting RFP (2) | Chloramphenicol | - | cgtttcaaagttcgtat |
| pCJH014 | Plasmid expressing Cas12c sgRNA targeting RFP (3) | Chloramphenicol | - | ggacatcctgtccccgc |
| pCJH015 | Plasmid expressing a Cas12c non-targeting sgRNA | Chloramphenicol | - | ttttttgtacttggtac |
| pCJH026 | Plasmid expressing a Cas12c sgRNA targeting GFP and another sgRNA targeting RFP expressed in one transcript for testing multiplexed repression | Chloramphenicol | Yes | TCCCAATTCTTGTTGAA; GCGAGTAGCGAAGACGT |
| pCJH029 | Plasmid expressing Cas12a guide RNA targeting the promoter of integrated GFP/RFP | Chloramphenicol | - | ACAGCTAGCTCAGTCCTAGGTATAATAG |
| pCJH030 | Plasmid expressing Cas12a guide RNA targeting the RBS sequence of integrated GFP/RFP | Chloramphenicol | - | ATGAATTCAGATCTATTATACCTAGGAC |
| pCJH031 | Plasmid expressing Cas12a guide RNA targeting GFP (1) | Chloramphenicol | Yes | ACTGGAGTTGTCCCAATTCTTGTTGAAT |
| pCJH032 | Plasmid expressing Cas12a guide RNA targeting GFP (2) | Chloramphenicol | - | TGCCCATTAACATCACCATCTAATTCAA |
| pCJH033 | Plasmid expressing Cas12a guide RNA targeting RFP (1) | Chloramphenicol | - | aaagttcgtatggaaggttccgttaacg |
| pCJH034 | Plasmid expressing Cas12a guide RNA targeting RFP (2) | Chloramphenicol | - | ATAACGTCTTCGCTACTCGCCATGGTAC |
| pCJH035 | Plasmid expressing a Cas12a non-targeting guide RNA | Chloramphenicol | - | ttttttgtacttggtacagcaggattca |
| pCJH037 | Plasmid expressing Cas12c sgRNA targeting dnaA (1) | Chloramphenicol | - | TCACTTTCGCTTTGGCA |
| pCJH038 | Plasmid expressing Cas12c sgRNA targeting dnaA (2) | Chloramphenicol | - | GCAGCAGTGTCTTGCCC |
| pCJH039 | Plasmid expressing Cas12c sgRNA targeting murD (1) | Chloramphenicol | - | GCTGATTATCAGGGTAA |
| pCJH040 | Plasmid expressing Cas12c sgRNA targeting murD (2) | Chloramphenicol | - | TCGTCATTATCGGCCTG |
| pCJH041 | Plasmid expressing Cas12c sgRNA targeting rplJ (1) | Chloramphenicol | - | GCTTTAAATCTTCAAGA |
| pCJH042 | Plasmid expressing Cas12c sgRNA targeting rplJ (2) | Chloramphenicol | - | TTGCTGAAGTCAGCGAA |
| pCJH043 | Plasmid expressing Cas12c sgRNA targeting rpoB (1) | Chloramphenicol | - | GTTTACTCCTATACCGA |
| pCJH044 | Plasmid expressing Cas12c sgRNA targeting rpoB (2) | Chloramphenicol | - | GTAAACGTCCACAAGTT |
| pCJH045 | Plasmid expressing Cas12a guide RNA targeting dnaA (1) | Chloramphenicol | - | GCTTTGGCAGCAGTGTCTTGCCCGATTG |
| pCJH046 | Plasmid expressing Cas12a guide RNA targeting dnaA (2) | Chloramphenicol | - | GCAGCAGTGTCTTGCCCGATTGCAGGAT |
| pCJH047 | Plasmid expressing Cas12a guide RNA targeting murD (1) | Chloramphenicol | - | CTGCGTGGACTTTTTCCTCGCTCGCGGT |
| pCJH048 | Plasmid expressing Cas12a guide RNA targeting murD (2) | Chloramphenicol | - | CTCGCTCGCGGTGTGACGCCGCGCGTTA |
| pCJH049 | Plasmid expressing Cas12a guide RNA targeting rplJ (1) | Chloramphenicol | - | AATCTTCAAGACAAACAAGCGATTGTTG |
| pCJH050 | Plasmid expressing Cas12a guide RNA targeting rplJ (2) | Chloramphenicol | - | GCTACTTCGCTGACTTCAGCAACAATCG |
| pCJH051 | Plasmid expressing Cas12a guide RNA targeting rpoB (1) | Chloramphenicol | - | CTCCTATACCGAGAAAAAACGTATTCGT |
| pCJH052 | Plasmid expressing Cas12a guide RNA targeting rpoB (2) | Chloramphenicol | - | GTAAACGTCCACAAGTTCTGGATGTACC |
| pCJH053 | Plasmid expressing Cas12c sgRNA targeting Lambda cro (1) | Chloramphenicol | - | gaacaacgcataaccct |
| pCJH054 | Plasmid expressing Cas12c sgRNA targeting Lambda cro (2) | Chloramphenicol | - | ggcaaaccaagacagct |
| pCJH055 | Plasmid expressing a Cas12c non-targeting sgRNA | Chloramphenicol | - | gctagcatgactggtgg |
| pCJH057 | Plasmid expressing Cas12a guide RNA targeting Lambda cro | Chloramphenicol | - | ggcaaaccaagacagctaaagatctcgg |
| pCJH058 | Plasmid expressing a Cas12a non-targeting guide RNA | Chloramphenicol | - | gcggtgaagtcctgactgcgttcgctcg |
| pCJH059 | Plasmid expressing Cas12c tracrRNA and a GFP-targeting pre-crRNA separately | Chloramphenicol | Yes | TCCCAATTCTTGTTGAA |
| pCJH061 | Plasmid expressing two Cas12c non-targeting sgRNAs connected in one transcript as a control for multiplexed repression | Chloramphenicol | - | ttttttgtacttggtac; tcgagtaaggatctcca |
| pCJH064 | Plasmid expressing Cas12c tracrRNA and a non-targeting pre-crRNA separately | Chloramphenicol | - | ttttttgtacttggtac |
| pCJH065 | Plasmid expressing Cas12c tracrRNA and GFP-targeting crRNA separately, with the crRNA ending with a HDV ribozyme sequence. | Chloramphenicol | Yes | TCCCAATTCTTGTTGAA |
| pCJH066 | Plasmid expressing Cas12c tracrRNA and non-targeting crRNA separately, with the crRNA ending with a HDV ribozyme sequence. | Chloramphenicol | - | ttttttgtacttggtac |

### Supplementary Table S2: Oligonucleotide sequences

**Table S2A: ssDNA oligonucleotides PAGE-purified for assays**

| ID | Description | Length | Sequence | Source |
|---|---|---|---|---|
| dCJH0073 | Cas12c Non-target Strand DNA | 53 | GCCTGCCCGCAGATTGatcaataccaaactctgGCGGCGTAAACTTTCCAGTC | IDT |
| dCJH0074 | Cas12c Target Strand DNA | 53 | GACTGGAAAGTTTACGCCGCcagagtttggtattgatCAATCTGCGGGCAGGC | IDT |
| dCJH0086 | 17-nt Mismatched Non-target Strand or Bubbled dsDNA | 53 | GCCTGCCCGCAGATTGttttttgtacttggtacGCGGCGTAAACTTTCCAGTC | IDT |
| dCJH0096 | Cas12c No PAM Non-target Strand DNA | 53 | GCCTGCCCGCAGATACatcaataccaaactctgGCGGCGTAAACTTTCCAGTC | IDT |
| dCJH0097 | Cas12c No PAM Target Strand DNA | 53 | GACTGGAAAGTTTACGCCGCcagagtttggtattgatGTATCTGCGGGCAGGC | IDT |
| dCJH0098 | Cas12c No Protopacer Target Strand DNA | 53 | GACTGGAAAGTTTACGCCGCgtaccaagtacaaaaaaCAATCTGCGGGCAGGC | IDT |

**Table S2B: ssDNA oligos used to generate the ssRNA using in vitro transcription (IVT)**

| ID | Description | Length | Sequence | Source |
|---|---|---|---|---|
| oCJH0017 | Used in a PCR reaction to generate the dsDNA template for IVT of rCJH007 | 58 | GGCTCTCCCTTAGCCATCCGAGTTTCCTCGGATGCCCAGGTCGGACCGCGAGGAGGTG | IDT |
| oCJH0031 | Used in a PCR reaction to generate the dsDNA template for IVT of rCJH006 | 47 | GTCGAAATTAATACGACTCACTATAGGTGGTATCTGATGAGGCCTTC | IDT |
| oCJH0032 | Used in a PCR reaction to generate the dsDNA template for IVT of rCJH006 | 43 | AGATTTCAGGTCGCTCTAGCTGCGCGCCCGGACATTGAGGTAC | IDT |
| oCJH0033 | Used in a PCR reaction to generate the dsDNA template for IVT of rCJH006 | 59 | GTATCTGATGAGGCCTTCGGGCCGAAACGGTGAAAGCCGTAATACCACCCGTGCATTTC | IDT |
| oCJH0034 | Used in a PCR reaction to generate the dsDNA template for IVT of rCJH006 | 53 | CCGGACATTGAGGTACGGATCATTGATCCAGAAATGCACGGGTGGTATTACGG | IDT |
| oCJH0035 | Used in PCR reactions to generate the dsDNA template for IVT of rCJH007, rCJH010 | 49 | GTCGAAATTAATACGACTCACTATAgggtaacgagcaggattcaggttg | IDT |
| oCJH0036 | Used in a PCR reaction to generate the dsDNA template for IVT of rCJH007 | 60 | aacgagcaggattcaggttgggtttgaggATCAATACCAAACTCTGagcaggatGGGTCG | IDT |
| oCJH0037 | Used in a PCR reaction to generate the dsDNA template for IVT of rCJH007 | 51 | GACCGCGAGGAGGTGGAGATGCCATGCCGACCCatcctgctCAGAGTTTGG | IDT |
| oCJH0042 | Used in a PCR reaction to generate the dsDNA template for IVT of rCJH010 | 42 | GGCTCTCCCTTAGCCATCCGAGTTTCCTCGGATGCCCAGGTC | IDT |
| oCJH0043 | Used in a PCR reaction to generate the dsDNA template for IVT of rCJH010 | 60 | taacgagcaggattcaggttgggtttgaggATCAATACCAAACTCTGGGGTCGGCATGGC | IDT |
| oCJH0051 | T7 promoter oligo in the forward direction for IVT of rCJH019, rCJH024, and rCJH040 | 17 | TAATACGACTCACTATA | IDT |
| oCJH0044 | Used in a PCR reaction to generate the dsDNA template for IVT of rCJH010 | 58 | TCGGATGCCCAGGTCGGACCGCGAGGAGGTGGAGATGCCATGCCGACCCCAGAGTTTG | IDT |
| oCJH0060 | Used in a PCR reaction to generate the dsDNA template for IVT of rCJH017 | 42 | GTCGAAATTAATACGACTCACTATAagcaggattcaggttgg | IDT |
| oCJH0061 | Used in PCR reactions to generate the dsDNA template for IVT of rCJH017, rCJH018 | 42 | GGCTCTCCCTTAGCCATCCGAGTTTCCTCGGATGCCCAGGTC | IDT |
| oCJH0062 | Used in a PCR reaction to generate the dsDNA template for IVT of rCJH017 | 60 | gcaggattcaggttgggtttgaggATCAATACCAAACTCTGagcaggatGGGTCGGCATG | IDT |
| oCJH0063 | Used in PCR reactions to generate the dsDNA template for IVT of rCJH017, rCJH018 | 53 | TCGGATGCCCAGGTCGGACCGCGAGGAGGTGGAGATGCCATGCCGACCCatcc | IDT |
| oCJH0064 | Used in a PCR reaction to generate the dsDNA template for IVT of rCJH018 | 60 | ATAggattcaggttgggtttgaggATCAATACCAAACTCTGagcaggatGGGTCGGCATG | IDT |
| oCJH0065 | Used in a PCR reaction to generate the dsDNA template for IVT of rCJH018 | 43 | GTCGAAATTAATACGACTCACTATAggattcaggttgggtttg | IDT |
| oCJH0066 | Used to anneal with the T7 promoter primer (oCJH0051) for IVT of rCJH019 | 99 | aacctgaatcctgctCAGAGTTTGGTATTGATCAGAGTTTGGTATTGATcctcaaacccaacctgaatcctgctcgttacccTATAGTGAGTCGTATTA | IDT |
| oCJH0071 | Used to anneal with the T7 promoter primer (oCJH0051) for IVT of rCJH024 | 155 | aacctgaatcctgctCAGAGTTTGGTATTGATcctcaaacccaacctgaatcctgctTTTCAGATTTCAGGTCGCTCTAGCTGCGCGCCCGGACATTGAGGTACGGATCATTGATCCAGAAATGCACGGGTGGTATCCTATAGTGAGTCGTATTA | IDT |
| oCJH0076 | Used in a PCR reaction to generate the dsDNA template for IVT of rCJH033 | 60 | ACTCACTATAGGCCAGTCCTGATGAGGCCTTCGGGCCGAAACGGTGAAAGCCGTAGACTG | IDT |
| oCJH0077 | Used in a PCR reaction to generate the dsDNA template for IVT of rCJH033 | 56 | TGAAAGCCGTAGACTGGAAAGTTTACGCCGCcagagtttggtattgatCAATCTGC | IDT |
| oCJH0078 | Used in a PCR reaction to generate the dsDNA template for IVT of rCJH033 | 52 | ggtattgatCAATCTGCGGGCAGGCGGGTCGGCATGGCATCTCCACCTCCTC | IDT |
| oCJH0079 | Used in a PCR reaction to generate the dsDNA template for IVT of rCJH033 | 50 | CATCTCCACCTCCTCGCGGTCCGACCTGGGCATCCGAGGAAACTCGGATG | IDT |
| oCJH0080 | Used in a PCR reaction to generate the dsDNA template for IVT of rCJH033 | 29 | GGCTCTCCCTTAGCCATCCGAGTTTCCTC | IDT |
| oCJH0081 | Used in a PCR reaction to generate the dsDNA template for IVT of rCJH033 | 31 | GTCGAAATTAATACGACTCACTATAGGCCAG | IDT |
| oCJH0087 | Used in PCR reactions to generate the dsDNA template for IVT of rCJH038, rCJH041 | 39 | GTCGAAATTAATACGACTCACTATAGGATACCACCCGTG | IDT |
| oCJH0088 | Used in PCR reactions to generate the dsDNA template for IVT of rCJH038, rCJH041 | 30 | GGCTCTCCCTTAGCCATCCGAGTTTCCTCG | IDT |
| oCJH0089 | Used in PCR reactions to generate the dsDNA template for IVT of rCJH038, rCJH041 | 42 | AGGATACCACCCGTGCATTTCTGGATCAATGATCCGTACCTC | IDT |
| oCJH0090 | Used in PCR reactions to generate the dsDNA template for IVT of rCJH038, rCJH041 | 56 | GATCAATGATCCGTACCTCAATGTCCGGGCGCGCAGCTAGAGCGACCTGAAATCTG | IDT |
| oCJH0091 | Used in a PCR reaction to generate the dsDNA template for IVT of rCJH038 | 44 | AGCGACCTGAAATCTGAAAagcaggattcaggttgggtttgagg | IDT |
| oCJH0092 | Used in PCR reactions to generate the dsDNA template for IVT of rCJH038, rCJH041 | 60 | caggttgggtttgaggATCAATACCAAACTCTGGGGTCGGCATGGCATCTCCACCTCCTC | IDT |
| oCJH0093 | Used in PCR reactions to generate the dsDNA template for IVT of rCJH038, rCJH041 | 51 | GCATCTCCACCTCCTCGCGGTCCGACCTGGGCATCCGAGGAAACTCGGATG | IDT |

| | | | | |
|---|---|---|---|---|
| oCJH0094 | Used to anneal with the T7 promoter primer (oCJH0051) for IVT of rCJH040 | 149 | aacctgaatcctgctCAGAGTTTGGTATTGATcctcaaacccaacctgaatTTTCAGATTTCAGGTCGCTCTAGCTGCGCGCCCGGACATTGAGGTACGGATCATTGATCCAGAAATGCACGGGTGGTATCCTATAGTGAGTCGTATTA | IDT |
| oCJH0095 | Used in a PCR reaction to generate the dsDNA template for IVT of rCJH041 | 38 | AGCGACCTGAAATCTGAAAattcaggttgggtttgagg | IDT |

**Table S2C: ssRNA oligos used in this study**

| ID | Description | Length | Sequence | Source |
|---|---|---|---|---|
| rCJH006 | Cas12c tracrRNA | 75 | AUACCACCCGUGCAUUUCUGGAUCAAUGAUCCGUACCUCAAUGUCCGGGCGCGCAGCUAGAGCGACCUGAAAUCU | PCR IVT |
| rCJH007 | Cas12c 5' extended pre-crRNA (8nt random + RS + 8nt of R) | 58 | GGGUAACGAGCAGGAUUCAGGUUGGGUUUGAGGAUCAAUACCAAACUCUG AGCAGGAU | PCR IVT |
| rCJH010 | Mature Cas12c crRNA (for rCJH007) (8nt random + RS) | 50 | GGGUAACGAGCAGGAUUCAGGUUGGGUUUGAGGAUCAAUACCAAACUCUG | PCR IVT |
| rCJH017 | Cas12c WT pre-crRNA (RS + 8nt of R) | 50 | AGCAGGAUUCAGGUUGGGUUUGAGGAUCAAUACCAAACUCUG AGCAGGAU | PCR IVT |
| rCJH018 | Cas12c pre-crRNA (21-nt shortened R + S + 8nt of R) | 46 | GGAUUCAGGUUGGGUUUGAGGAUCAAUACCAAACUCUG AGCAGGAU | PCR IVT |
| rCJH019 | Cas12c double spacer pre-crRNA (8nt random + RSS + 15nt of R) | 82 | GGGUAACGAGCAGGAUUCAGGUUGGGUUUGAGGAUCAAUACCAAACUCUG AUCAAUACCAAACUCUGAGCAGG AUUCAGGUU | IDT IVT |
| rCJH024 | Cas12c pre-sgRNA version 1 (tracrRNA + GAAA + RS + 15nt R) | 138 | GGAUACCACCCGUGCAUUUCUGGAUCAAUGAUCCGUACCUCAAUGUCCGGGCGCGCAGCUAGAGCGACCUGAAAUCUGAAAAGCAGGAUUCAGGUUGGGUUUGAGGAUCAAUACCAAACUCUG AGCAGGAUUCAGGUU | IDT IVT |
| rCJH026 | 3' 6-FAM labeled Cas12c pre-crRNA (RS + 11nt of R) | 49 | GGAUUCAGGUUGGGUUUGAGGAUCAAUACCAAACUCUG AGCAGGAUUCA/36-FAM/ | IDT |
| rCJH027 | 3' 6-FAM labeled 11nt R Cas12c with 5' P | 11 | /5Phos/AGCAGGAUUCA/36-FAM/ | IDT |
| rCJH028 | 3' 6-FAM labeled 11nt R Cas12c with 5' OH | 11 | AGCAGGAUUCA/36-FAM/ | IDT |
| rCJH029 | Multi-phosphorothioated rCJH018 Cas12c pre-crRNA (21-nt shortened R + S + 8nt of R) | 46 | GGAUUCAGGUUGGGUUUGAGGAUCAAUACCAAACUCU*G*A*G*CAGGAU | IDT |
| rCJH030 | Phosphorothioated rCJH018 Cas12c pre-crRNA (21-nt shortened R + S + 8nt of R) | 46 | GGAUUCAGGUUGGGUUUGAGGAUCAAUACCAAACUCU*G AGCAGGAU | IDT |
| rCJH031 | Phosphorothioated rCJH018 Cas12c pre-crRNA (21-nt shortened R + S + 8nt of R) | 46 | GGAUUCAGGUUGGGUUUGAGGAUCAAUACCAAACUCUG*AGCAGGAU | IDT |
| rCJH032 | Phosphorothioated rCJH018 Cas12c pre-crRNA (21-nt shortened R + S + 8nt of R) | 46 | GGAUUCAGGUUGGGUUUGAGGAUCAAUACCAAACUCUG A*GCAGGAU | IDT |
| rCJH033 | (ssRNA) Cas12c target strand (cleaved by HHrz and HDVrz) | 53 | GACUGGAAAGUUUACGCCGCCAGAGUUUGGUAUUGAUCAAUCUGCGGGCAGGC | PCR IVT |
| rCJH037 | Mature Cas12c crRNA (for rCJH018) (21-nt shortened R + S) | 38 | GGAUUCAGGUUGGGUUUGAGGAUCAAUACCAAACUCUG | IDT |
| rCJH038 | Mature sgRNA version 1 (processed version of rCJH024) | 123 | GGAUACCACCCGUGCAUUUCUGGAUCAAUGAUCCGUACCUCAAUGUCCGGGCGCGCAGCUAGAGCGACCUGAAAUCUGAAAAGCAGGAUUCAGGUUGGGUUUGAGGAUCAAUACCAAACUCUG | PCR IVT |
| rCJH039 | Partial DNA version of rCJH018 Cas12c pre-crRNA (21-nt shortened R + S + 8nt of R) | 46 | rGrGrArUrUrCrArGrGrUrUrGrGrGrUrUrUrGrArGrGrArUrCrArArUrArCrCrArArArCrUrC(d)T(d)G(d)A(d)GrCrArGrGrArU | IDT |
| rCJH040 | Cas12c sgRNA version 2 (tracrRNA + GAAA + 19nt R+ S + 15nt R) | 132 | GGAUACCACCCGUGCAUUUCUGGAUCAAUGAUCCGUACCUCAAUGUCCGGGCGCGCAGCUAGAGCGACCUGAAAUCUGAAAAUUCAGGUUGGGUUUGAGGAUCAAUACCAAACUCUG AGCAGGAUUCAGGUU | IDT IVT |
| rCJH041 | Mature sgRNA version 2 (processed version of rCJH040) | 117 | GGAUACCACCCGUGCAUUUCUGGAUCAAUGAUCCGUACCUCAAUGUCCGGGCGCGCAGCUAGAGCGACCUGAAAUCUGAAAAUUCAGGUUGGGUUUGAGGAUCAAUACCAAACUCUG | PCR IVT |
| rCJH042 | Partial DNA version of rCJH018 Cas12c pre-crRNA (21-nt shortened R + S + 8nt of R) | 46 | rGrGrArUrUrCrArGrGrUrUrGrGrGrUrUrUrGrArGrGrArUrCrArArUrArCrCrArArArCrUrC(d)TrGrArGrCrArGrGrArU | IDT |
| rCJH043 | Partial DNA version of rCJH018 Cas12c pre-crRNA (21-nt shortened R + S + 8nt of R) | 46 | rGrGrArUrUrCrArGrGrUrUrGrGrGrUrUrUrGrArGrGrArUrCrArArUrArCrCrArArArCrUrCrU(d)GrArGrCrArGrGrArU | IDT |
| rCJH044 | Partial DNA version of rCJH018 Cas12c pre-crRNA (21-nt shortened R + S + 8nt of R) | 46 | rGrGrArUrUrCrArGrGrUrUrGrGrGrUrUrUrGrArGrGrArUrCrArArUrArCrCrArArArCrUrCrUrG(d)ArGrCrArGrGrArU | IDT |
| rCJH045 | Partial DNA version of rCJH018 Cas12c pre-crRNA (21-nt shortened R + S + 8nt of R) | 46 | rGrGrArUrUrCrArGrGrUrUrGrGrGrUrUrUrGrArGrGrArUrCrArArUrArCrCrArArArCrUrCrUrGrA(d)GrCrArGrGrArU | IDT |

Notes:

IDT IVT: A long ssDNA oligo from IDT were directly annealed with a T7 promoter oligo (oCJH0051) and transcribed

PCR IVT: multiple IDT oligos were used in a PCR reaction to create the dsDNA template with a T7 promoter sequence for IVT, on which IVT was then performed

IDT: Oligos ordered from Integrated DNA Technologies

R = Repeat sequence; S = Spacer sequence

**Table S2D: DNA oligos used to assemble *in vitro* transcription templates**

| oligo ID | IVT template assembled by PCR or direct annealing from DNA oligos: | ribozyme(s) in initial transcript? |
|---|---|---|
| rCJH006 | oCJH0031, oCJH0032, oCJH0033, oCJH0034 | HHrz |
| rCJH007 | oCJH0035, oCJH0017, oCJH0036, oCJH0037 | HDVrz |
| rCJH010 | oCJH0035, oCJH0042, oCJH0043, oCJH0044 | HDVrz |
| rCJH017 | oCJH0060, oCJH0061, oCJH0062, oCJH0063 | HDVrz |
| rCJH018 | oCJH0065, oCJH0061, oCJH0064, oCJH0063 | HDVrz |
| rCJH019 | oCJH_0066, oCJH0051_T7oligo | None |
| rCJH024 | oCJH_0071, oCJH0051_T7oligo | None |
| rCJH033 | oCJH0080, oCJH0081, oCJH0076, oCJH0077, oCJH0078, oCJH0079 | HHrz, HDVrz |
| rCJH038 | oCJH0087, oCJH0088, oCJH0089, oCJH0090, oCJH0091, oCJH0092, oCJH0093 | HDVrz |
| rCJH040 | oCJH_0094, oCJH0051_T7oligo | None |
| rCJH041 | oCJH0087, oCJH0088, oCJH0089, oCJH0090, oCJH0095, oCJH0092, oCJH0093 | HDVrz |

Key:

HHrz: hammerhead ribozyme on 5′ end of transcript

HDVrz: hepatitis delta virus ribozyme on 3′ end of transcript

### Supplementary Table S3: Plasmids and oligonucleotides used in experiments

| Table S3A: Plasmid or oligo ID used in main-text figures | |
|---|---|
| | **Plasmid or oligo ID** |
| **Figure 1B** | tracrRNA: rCJH006<br>crRNA: rCJH010<br>dsDNA (TS/NTS): dCJH0074+dCJH0073<br>Bubbled dsDNA (TS/bubbledNTS): dCJH0074+dCJH0086<br>ssDNA: dCJH0074<br>ssRNA: rCJH033 |
| **Figure 2A** | pre-crRNA: rCJH018, tracrRNA: rCJH006<br>RNA marker: rCJH037 |
| **Figure 2C** | pre-crRNA: rCJH026<br>tracrRNA: rCJH006<br>RNA markers: rCJH027 and rCJH028 |
| **Figure 2D** | pre-crRNA: rCJH039<br>tracrRNA: rCJH006<br>RNA marker: rCJH037 |
| **Figure 2E** | Pre-crRNA: rCJH042, rCJH043, rCJH044, rCJH045<br>tracrRNA: rCJH006<br>RNA Marker: rCJH037 |
| **Figure 3B** | pre-sgRNA: rCJH024 and rCJH040<br>RNA markers: rCJH038 and rCJH041 |
| **Figure 4A** | tracrRNA: rCJH006<br>crRNA: rCJH010<br>dsDNA (TS/NTS): dCJH0074+dCJH0073<br>Bubbled dsDNA (TS/bubbledNTS): dCJH0074+dCJH0086<br>dsDNA with no PAM (NTS/TS): dCJH0096+dCJH0097<br>Non-complementary dsDNA (NTS/TS): dCJH0086+dCJH0098<br>ssDNA: dCJH0074<br>ssRNA: rCJH033 |
| **Figure 4B** | tracrRNA: rCJH006<br>pre-crRNA: rCJH018<br>Mature crRNA: rCJH037<br>Phosphorothioated pre-crRNA: rCJH029<br>dsDNA (5' radiolabeled NTS/TS): dCJH0073+dCJH0074 |
| **Figure 4C** | tracrRNA: rCJH006<br>pre-crRNA: rCJH018<br>Mature crRNA: rCJH037<br>dsDNA (5' radiolabeled NTS/TS): dCJH0073+dCJH0074 |
| **Figure 5B** | Cas plasmids: pCJH027 or pCJH028<br>Guide RNA plasmids: pCJH029, pCJH030, pCJH031, pCJH032, pCJH033, pCJH0034, and pCJH035 |
| **Figure 5C** | Cas plasmids: pCJH019 or pCJH020<br>sgRNA plasmids: pCJH007, pCJH008, pCJH009, pCJH010, pCJH011, pCJH012, pCJH013, pCJH014, and pCJH015 |
| **Figure 5D&E** | Cas plasmids: pCJH021 or pCJH022<br>sgRNA plasmids: pCJH009 and pCJH015<br>Dual RNA guide plasmids: pCJH059 and pCJH064<br>Dual RNA with HDVrz guide plasmids: pCJH065 and pCJH066 |
| **Figure 5F&G** | Cas plasmids: pCJH019 or pCJH020<br>sgRNA plasmids: pCJH009 and pCJH015<br>Dual RNA guide plasmids: pCJH059 and pCJH064<br>Dual RNA with HDVrz guide plasmids: pCJH065 and pCJH066 |
| **Figure 5H** | Cas plasmid: pCJH019<br>sgRNA plasmids: pCJH026 and pCJH061 |
| **Figure 6A** | Cas12c plasmid: pCJH019<br>Cas12c sgRNA plasmids: pCJH053, pCJH054, pCJH055<br>Cas12a plasmids: pCJH027 and pCJH028<br>Cas12a guide plasmids: pCJH057 and pCJH058 |

| Table S3B: Plasmid or oligo ID used in supplementary figures | |
|---|---|
| | **Plasmid or oligo ID** |
| **Figure S1A** | Pre-crRNA: rCJH007, rCJH017, rCJH018, and rCJH019<br>tracrRNA: rCJH006<br>RNA marker: rCJH010 |
| **Figure S1B** | pre-crRNA: rCJH026<br>tracrRNA: rCJH006 |
| **Figure S1C** | Pre-crRNA: rCJH018, rCJH029, rCJH030, rCJH031, rCJH032<br>tracrRNA: rCJH006<br>RNA marker: rCJH037 |
| **Figure S1D** | Pre-crRNA: rCJH018 and rCJH029<br>tracrRNA: rCJH006 |
| **Figure S2A** | pre-crRNA: rCJH017<br>tracrRNA: rCJH006 |
| **Figure S2B** | pre-crRNA: rCJH017<br>tracrRNA: rCJH006 |
| **Figure S3A** | Cas plasmids: pCJH027 or pCJH028<br>Guide RNA plasmids: pCJH029, pCJH030, pCJH031, pCJH032, pCJH033, pCJH0034, and pCJH035 |
| **Figure S3B** | Cas plasmids: pCJH019 or pCJH020<br>sgRNA plasmids: pCJH007, pCJH008, pCJH009, pCJH010, pCJH011, pCJH012, pCJH013, pCJH014, and pCJH015 |
| **Figure S3C** | Cas plasmids: pCJH027 or pCJH028<br>Guide RNA plasmids: pCJH045, pCJH046, pCJH047, pCJH048, pCJH049, pCJH050, pCJH051, pCJH052, pCJH035 |
| **Figure S3D** | Cas plasmid: pCJH019<br>sgRNA plasmids: pCJH037, pCJH038, pCJH039, pCJH040, pCJH041, pCJH042, pCJH043, pCJH044, pCJH015 |
| **Figure S3E** | pre-crRNA: rCJH017<br>tracrRNA: rCJH006 |
